# What is common to memory retrieval and relational reasoning? Testing neural overlap in anterior lateral prefrontal cortex

**DOI:** 10.64898/2026.09.06.749563

**Authors:** Allison Chen, Maggie Vashel, Mariam Aly, Silvia A. Bunge

**Author notes:** Corresponding author: Allison Chen, Interdepartmental Neuroscience Program University of California, Irvine, Irvine, CA 92697.

## Abstract

Anterior lateral prefrontal cortex (aLPFC) has been consistently implicated in retrieval monitoring and relational reasoning. However, it is unknown whether aLPFC supports a cognitive process that is common to memory retrieval and reasoning tasks. To inform this question, a first step is to ascertain whether there is, within individuals, spatial overlap in aLPFC between tasks. A prior study found overlap in activation of left aLPFC between verbal analogy and source memory retrieval tasks, using tightly matched word stimuli (Westphal et al., 2016). To assess whether this overlap in activation extends to other stimulus types and task designs, and another sensory modality, we collected fMRI data from 35 healthy young adults as they completed an associative memory retrieval task along with four variants of a relational reasoning paradigm that reliably engages left aLPFC. The reasoning tasks followed a 2 (visual/auditory modality) x 2 (spatial/featural relations) x 2 (low/high relational complexity) design. In the associative memory task, participants studied word pairs and decided whether the two words were related. At retrieval, they viewed pairs of previously seen words and judged whether the pair was Intact (words paired together at encoding) or Recombined (previously seen words recombined into new pairs). Response times were slower for Recombined vs. Intact pairs, consistent with heightened retrieval monitoring. As predicted, aLPFC was more active for Recombined than Intact pairs, as well as for the manipulation of relational complexity across the four reasoning tasks. A group-level conjunction analysis demonstrated substantial overlap between memory and reasoning manipulations in left aLPFC (Glasser parcels a47r, p47r, a10p, and p10p). Within individuals, clusters defined from the reasoning task were also sensitive to retrieval monitoring demands. Further, there was within-participant overlap in the pattern of active vertices, as measured via Dice coefficients. Bayes factor testing within each parcel of interest demonstrated greater vertex-level overlap than expected by chance, particularly in p47r. Importantly, degree of overlap varied widely among individuals, and exploratory results showed that both cluster-level activation and vertex overlap in p47r correlated with memory discriminability (d’). Thus, we hypothesize that neural overlap in this area reflects engagement in relational thinking during memory retrieval monitoring. Together, these results move beyond general characterizations of aLPFC function by testing whether it meets a precondition—individual-level neural overlap across varied tasks—of a unified neurocognitive function in support of reasoning and memory retrieval.

## 1 INTRODUCTION

Within human prefrontal cortex (PFC), the most anterior aspect is a large, heterogeneous area whose functional organization and mechanistic contributions to cognition are of abiding interest but are not well understood. One of the key challenges to understanding its functions is that there is no adequate homolog in other species (Vendetti & Bunge, 2014; Rosa et al., 2019). Anterior PFC in humans, relative to non-human primates, is disproportionately large (Semendeferi et al., 2011; Donahue et al., 2018), more anatomically variegated (Rosa et al., 2019), and has different patterns of white matter projections (Vendetti & Bunge, 2014), functional connectivity (Mars et al., 2011), and cellular morphology (Semendeferi et al., 2011). Despite these species differences, theories of its function often combine findings from human and non-human primate species.

Another challenge is the inconsistent use of nomenclature in this literature. Anterior PFC is often referred to interchangeably as Brodmann’s area (BA) 10. As one of the largest areas defined in Brodmann’s cytoarchitectonic atlas, however, human BA 10 is anatomically and functionally dissociable into lateral, medial, and orbital portions (Gilbert et al., 2005; Liu et al., 2013), with the clearest dissociation between lateral and medial portions (Burgess et al., 2003; Gilbert et al., 2010; Bahlmann et al., 2015; Moayedi et al., 2015; Altmayer et al., 2026). In fact, even when restricting focus to the most polar portion of PFC, one finds anatomical dissociations (Bludau et al., 2013; Orr et al., 2015). Within lateral PFC (LPFC), the anterior portion (aLPFC) is variably called the lateral frontal pole, rostrolateral PFC (RLPFC), or anterior inferior frontal gyrus — or more generally as BA 10, frontopolar cortex, anterior PFC, or rostral PFC, despite the aforementioned heterogeneity of anterior PFC. Moreover, activations that are reported as residing in lateral BA 10 often traverse boundaries (10/46, 10/47) or may even be localized entirely to BA 46 or 47 (Bunge et al., 2004; Bunge et al., 2005; Cho et al., 2010; Westphal et al., 2016).

Beyond the issue of inconsistent anatomical definitions and nomenclature, aLPFC has been linked to a variety of higher-order cognitive functions (Christoff & Gabrieli, 2000; Ramnani & Owen, 2004; Koechlin & Hyafil, 2007). Patients with neurological insults limited to or including aLPFC exhibit difficulties with complex cognitive tasks such as prospective memory (Burgess et al., 2003) and multi-tasking (Burgess et al., 2000; Dreher et al., 2008; Roca et al., 2011). Patients with damage to this region also exhibit increased confabulation, as well as greater difficulty with source memory retrieval (Duarte et al., 2005; Turner et al., 2008; Blumenfeld & Ranganath, 2019). Additionally, some studies have reported impaired performance on tests of reasoning (Roca et al., 2010; Woolgar et al., 2010; Urbanski et al., 2016; but see Burgess, 2000; Tranel et al., 2008).

fMRI and transcranial stimulation studies have linked aLPFC to a broader variety of cognitive domains, including memory retrieval monitoring and relational reasoning (Christoff & Gabrieli, 2000), as discussed below, and much more (Bunge 2004; Badre & D’Esposito, 2007; Boorman et al., 2009; Fleming & Dolan, 2012; Desrochers et al., 2015; Fandakova et al., 2018; Shekhar & Rahnev, 2018; Hogeveen et al., 2022). Despite its broad engagement, aLPFC activation is not elicited by all cognitively demanding tasks, and is not considered a core node of the Multiple Demand System (MDS) (Assem et al., 2020). Moreover, its activation does not scale with working memory load and cannot be explained by task difficulty per se (Burgess et al., 2003; Wendelken et al., 2008a; Wendelken et al., 2008b). Thus, with careful task manipulations, its activation profiles can be dissociated from those of more caudal LPFC regions.

In the domain of memory, numerous fMRI studies have reported aLPFC activation in episodic memory retrieval (e.g., Henson et al., 1999; Cabeza & Nyberg, 2000; Ranganath et al., 2000; Ranganath & Paller, 2000; Dobbins et al., 2002; Wagner et al., 2005; Ranganath et al., 2006; Simons et al., 2008). Moreover, one study showed improved episodic source memory retrieval after transcranial direct current stimulation to left aLPFC (Westphal et al., 2019). Activation in this general area is often observed on tasks designed to elicit recollection by requiring participants to resolve interference from highly similar lures (Ranganath et al., 2000), remember specific item-item associations in tests of associative memory or item-context associations in tests of source memory, or engage in free recall (Christoff & Gabrieli, 2000; Blumenfeld & Ranganath, 2019). Additionally, right aLPFC has been reported for manipulations of single-word recognition (e.g., Buckner 1998; Henson et al., 1999).

Integrating across many studies, memory researchers have arrived at the conclusion that aLPFC is linked neither to retrieval success nor effort *per se*, but rather to non-mnemonic aspects of memory judgments (e.g., Buckner et al., 1998; Rugg & Henson, 2002). One hypothesis that has been put forth is that aLPFC, perhaps particularly in the right hemisphere, is associated with a retrieval mode: an orientation towards cue-based retrieval (Nyberg et al., 1995; Lepage et al., 2000; Velanova et al., 2003; Underwood et al., 2015). Another is that aLPFC operates on the products of memory retrieval, for example in the form of post-retrieval monitoring in the service of accurate episodic memory (Ranganath et al., 2000; Dobbins et al., 2002; Rugg & Henson, 2002; Westphal et al., 2016).

In parallel, aLPFC activation, particularly in the left hemisphere, has been reported across many fMRI studies on reasoning, as measured with matrix reasoning, propositional analogy, and transitive inference tasks (e.g., Christoff et al., 2001; Kroger et al., 2002; Bunge et al., 2005; Wendelken & Bunge, 2010; Knowlton et al., 2012; Krawzcyk 2012; Vendetti & Bunge, 2014; Hobeika et al., 2016; Holyoak & Monti, 2021). Relational thinking, or the process of considering relations between active mental representations (Halford et al., 2010; Alexander, 2016), is central to such reasoning tasks. We have argued that it should be considered an executive function: that is, a mid-level control process that relies on attention and working memory and in turn supports higher-level cognitive abilities (Starr et al., 2023).

Consistent with research on episodic memory, fMRI studies on analogical reasoning suggest that aLPFC does not retrieve relations from semantic memory, but rather compares them after retrieval (Bunge et al., 2005; Bendetowicz et al., 2018). aLPFC is sensitive to relational complexity, and is therefore posited to facilitate the representation of higher-order relational structures – i.e., the relation between relations, such as in a propositional analogy (e.g., the brain is to mental processes as the small intestine is to digestion) or the reasoning paradigm used in this study.

A number of researchers have sought to integrate a constellation of these and other findings into a singular account of aLPFC function. In recent years, these accounts have largely converged around several closely related ideas. One set of theories—drawing on the observation that damage to aLPFC is associated with deficits in prospective memory, planning, and multi- tasking—relates to goal management (Koechlin et al., 1999; Braver & Bongiolotti, 2002; Ramnani & Owen, 2004; Mansouri et al., 2017). Another overarching theory is that aLPFC is involved in evaluating internally generated information, as needed when making inferences or retrieving information from memory (Christoff et al., 2003), switching between internally and externally generated information (Burgess et al., 2007), or making metacognitive judgments (Fleming et al., 2012; Baird et al., 2013; Fleming et al., 2014; Morales et al., 2018; Shekhar & Rahnev, 2018). Yet another prevalent set of ideas situates aLPFC within a caudal-rostral hierarchy of function across LPFC (Fuster, 2001), such as a progression of levels of abstraction of representations across LPFC (Christoff et al., 2009) or a hierarchy of behavioral control (Fuster, 2004; Badre & D’Esposito, 2007), e.g., whereby more rostral regions represent increasingly hierarchically nested rule structures (Koechlin et al., 2003; Bunge & Zelazo, 2006; Azuar et al., 2014).

Some of these accounts may be overly complex, given the sheer simplicity of some of the tasks that engage aLPFC (Dobbins & Han, 2006; Wendelken et al., 2008). One simpler account, centered on relational thinking, is that aLPFC contributes whenever task performance requires the comparison or integration of several active mental representations; these representations may be any combination of concepts, memories, rules, possible actions, criteria, choices, and/or outcomes. With regards to episodic memory retrieval, for example, active comparison is needed when judging whether details of an event, such as which items were paired together, match a memory of a prior event , or when discriminating between highly similar targets and lures. With regards to reasoning paradigms, active comparison is needed to consider relations among items or sets of items. In other words, the relational thinking hypothesis holds that aLPFC helps to meet demands for active comparison of working memory representations, which are common to episodic memory retrieval and reasoning, along with other higher-level cognitive functions like planning and decision-making (Christoff & Gabrieli, 2000; Bunge & Wendelken, 2009).

Crucially, it is not merely a question of which one these closely related accounts is most apt, but rather whether any unifying account of aLPFC function is possible. Indeed, any claim of a common functional role across tasks presupposes that the same area is recruited across domains. However, this is not a given. At a macroscopic level, possible hemispheric asymmetries have been observed across a number of fMRI and transcranial stimulation studies, with preferential engagement of left aLPFC in studies of relational reasoning (e.g., Bunge et al., 2009) and retrieval monitoring (Ranganath et al., 2000; Dobbins et al., 2002; Westphal et al., 2016), and right aLPFC in metacognition (e.g., Fleming & Dolan, 2012), and retrieval mode (e.g., Nyberg et al., 1995; Lepage et al., 2000). This pattern of results hints at a possible functional dissociation between left and right aLPFC, which warrants careful consideration. At a more granular level, much of the existing theorizing has relied on qualitative comparisons of peak coordinates of group-averaged activations normalized to a standard template. This approach ignores the pronounced heterogeneity across individuals in the location and extent of activation, coupled with a lack of anatomical landmarks that could enable the direct comparison of activation foci across studies. Thus, the formulation of unifying hypotheses has been premature; as a first step, quantifying overlap in activation must be done at the single-subject level.

Direct empirical tests of cross-task overlap in aLPFC within individuals remain rare. A notable exception is a study by Westphal et al. (2016) that examined source memory and verbal analogical reasoning within the same sample. Using closely matched task structures, stimuli, and timing, participants encoded individual words under self-or other-referential judgments and later completed either a source memory task (identifying whether any word in a four-word array had been encoded under a given source) or an analogy task (judging whether two word pairs formed a valid analogy). Group-level univariate analyses revealed extensive overlap in activation throughout much of left LPFC, including aLPFC, in addition to some activation in right LPFC. Notably, a pattern classifier trained within participants to discriminate memory and reasoning trials pinpointed an area of overlap in left aLPFC, described as posterior RLPFC, as the region with the highest task decoding accuracy. These findings provide preliminary support for a shared substrate, but leave open questions regarding the precise spatial extent and functional specificity of overlap.

In contrast to the extensive group-level overlap observed in left aLPFC by Westphal and colleagues (2016), a study by Reynolds et al. (2006) found no such overlap for episodic memory and relational integration manipulations in a different paradigm, with a different type of memory manipulation. At encoding in that study, participants either judged separately whether each of two words in a pair represented an abstract or concrete concept (low integration demand), or whether the two words matched on this dimension, e.g., both abstract (high integration demand). Then, in a recognition test, they either judged separately whether each of two words was old or new (low integration demand), or whether the two words matched on this dimension, e.g., both old (high integration demand). The investigators found a region in right aLPFC that was sensitive both to memory retrieval demands (retrieval > encoding) and relational integration (high > low); by contrast, they found distinct foci for the two tasks in left aLPFC. However, their memory measure consisted of a comparison of retrieval vs encoding, and not of memory monitoring demands. Further, because the regions of interest (ROIs) were defined at the group level, as with Westphal et al. (2016), it is unknown whether individual participants showed task overlap in this region.

The present study extends Westphal et al. (2016), broadening the scope to include five additional memory and reasoning tasks: an associative memory retrieval task involving word pairs, alongside a set of four relational reasoning paradigms involving both spatial and featural (non-spatial) visual and auditory stimuli. By precisely characterizing neural overlap within individuals, we take a step towards refining our understanding of the role of aLPFC in two of the cognitive functions aLPFC has been most consistently implicated in: memory retrieval monitoring and relational reasoning. Of course, overlapping activation measured at the spatial scale of fMRI is insufficient to establish a shared neural substrate; however, minimal overlap would place important constraints on unified accounts of aLPFC function.

### 1.1 Approach

Our memory task consisted of an associative memory paradigm originally developed to examine relational luring (Ichien et al., 2023). In this task, participants studied pairs of words and later judged whether test pairs were Intact (studied together) or Recombined (studied separately). Prior behavioral work (Van Petten et al., 2002; Jou 2010; Ichien et al., 2023) showed that responses were faster and more accurate for Intact than Recombined trials, likely reflecting veridical memory for previously encountered pairings over and above the recognition of the individual words. Building on prior work (Van Petten et al., 2002; Achim & Lepage 2005; Horne et al. 2021), we operate under the assumption that post-retrieval monitoring demands are greater for Recombined than Intact trials. Compared to Intact pairs, correct performance for Recombined pairs should be associated with more retrieval monitoring both because (i) the familiarity signal from the individual items has to be discounted and (ii) because recombined pairs may trigger retrieval of one or both originally paired items.

Our reasoning task consisted of a composite of four relational matching tasks collected with the same participants, which required identifying similarities of relations across pairs of stimuli. We adopted a paradigm previously shown to reliably engage aLPFC (Christoff et al., 2003; Wendelken et al., 2011). The four tasks were identical in structure but varied in stimulus modality and the type of relational judgment required: visual stimuli involving semantic or spatial content, and auditory stimuli involving featural or spatial content (see preregistration: osf.io/fhjt2). Here we compare the memory task with a composite of all four Reasoning tasks. In the Supplement, we additionally present the same set of analyses for the visual-semantic (“Vis-What”) Reasoning task alone, as it was most similar to the memory paradigm in terms of the amount of data collected and type of relations involved. Indeed, while the Vis-What task featured visually presented objects as opposed to words, both required semantic judgments. In another study, we have performed direct comparisons across the Reasoning tasks (Vashel et al., in prep; preregistered at <u>osf.io/fhjt2</u>).

We conducted both group-and individual-level analyses to characterize task overlap. As an initial step, we computed whole-brain group-level conjunction and disjunction maps to identify common and distinct regions of activation across the memory and reasoning paradigms. These analyses allowed us to determine whether overlapping activation emerged at the group level, and in which aLPFC parcels. We then conducted individual-level analyses to more directly assess the degree of overlap between tasks within aLPFC. Specifically, we examined activation patterns within predefined aLPFC parcels to determine whether the same subregions were engaged across tasks.

Although not preregistered, our intention was to focus on the left hemisphere, for several reasons. First, our decision was informed by evidence from Neurosynth meta-analytic fMRI maps for the keywords “relational” and “retrieval” (Yarkoni et al., 2011), and on individual empirical studies showing a left-hemisphere bias for relational thinking – even with visuospatial, non-linguistic stimuli (e.g., Bunge et al., 2009). Second, we sought to expand on the prior study showing overlapping activation in left aLPFC for memory retrieval and reasoning within individuals (Westphal et al., 2016), in which both tasks involved linguistic stimuli (words). We report exploratory results for the right hemisphere in the Supplement, for which we had no clear predictions.

To characterize activation within aLPFC, we used the multimodal cortical parcellation scheme developed by Glasser et al. (2016), which enabled us to conduct analyses in native cortical surface space and then aggregate results across participants without relying on voxel coordinates in standardized template space. Use of the Glasser atlas enables consideration of results in light of other studies that have characterized activation profiles in aLPFC parcels (e.g., Assem et al., 2020). This atlas defines cortical areas based on several functional and structural metrics, including resting-state functional connectivity, cortical thickness, myelin mapping, and structural connectivity (Glasser et al., 2016). Although the Glasser parcellation incorporates activation patterns from seven fMRI tasks, including a relational matching task, task activation contributed relatively little weight in defining parcel boundaries compared with other structural and connectivity features. Thus, we did not expect strong correspondence between Glasser parcels and activation in this study.

We next examined whether clusters in aLPFC parcels identified at the individual-subject level from the Reasoning task were also sensitive to monitoring demands during memory retrieval, particularly in contrasts isolating activation related to a relational complexity manipulation. Because individuals were expected to differ in aLPFC activation, either due to strategic variability or limited signal-to-noise, we anticipated that not all participants would show detectable activation in this region. We therefore quantified the proportion of participants exhibiting task-related activation in a given aLPFC parcel and extracted clusters for subsequent analyses from those individuals who did.

To examine overlap at the vertex-level, we also tested whether the vertices most strongly implicated in the Reasoning task were also those most strongly engaged during memory retrieval. Using the top 5% of active vertices in each task, we quantified overlap by calculating Dice coefficients in each parcel for each participant, and then used Bayesian analyses to compare the degree of overlap across parcels.

We additionally conducted exploratory analyses examining whether level of activation in reasoning-related clusters was related to memory performance across individuals. In addition, we assessed overlap between tasks at a more granular level by testing for relationships between memory performance and patterns of activation across individual surface vertices.

To provide benchmarks for interpreting these results, we included two comparison regions. First, we examined primary visual cortex (V1), which is strongly stimulus-driven and therefore expected to show task-specific activation patterns. Second, we examined a region in dorsolateral PFC (DLPFC) that has been included in the MDS (Assem et al., 2020), a set of regions that is engaged across a wide variety of cognitively demanding tasks. These comparison regions allowed us to situate aLPFC overlap in relation to both stimulus-driven and putatively domain-general control regions.

### 1.2 Predictions

We predicted that memory retrieval and relational reasoning tasks would show overlapping activation within left aLPFC parcels, replicating and extending Westphal et al. (2016). However, several alternative outcomes are equally plausible. First, because this was the first fMRI study to employ the present retrieval paradigm, it was possible that the task would not reliably engage aLPFC. Second, even if the memory task were to recruit aLPFC, its activation patterns might not overlap with those for relational reasoning. Several prior studies have reported coordinates that correspond to parcel a47r in relational integration (Bunge et al., 2004; Bunge et al., 2005; Cho et al., 2010), while other studies have implicated parcel 45 in controlled semantic retrieval (Bunge et al., 2004; Souza et al., 2011). Thus, a single aLPFC subregion may not support relational comparison processes across both memory and reasoning paradigms.

## 2 METHODS

### 2.1 Participants

Participants were affiliates of UC Berkeley or members of the surrounding community, recruited as part of a larger relational reasoning study. Individuals enrolled in the study were fluent or native English speakers with normal or corrected-to-normal vision, normal hearing and motor capacity, and no history of neurological or major mental health disorders. Informed consent was obtained following procedures approved by the Institutional Review Board at UC Berkeley, and participants received either monetary compensation or course credit in an undergraduate Psychology class.

Fifty participants completed fMRI scans of both a relational reasoning task and a memory retrieval task. Following criteria set for the Reasoning task, three participants were excluded for poor behavioral performance, and five participants were excluded because of excessive head motion (see Section 2.5 for all exclusion criteria). Following criteria set for the memory task, one participant was excluded due to issues recording their behavioral responses and six were excluded because of excessive head motion. The final sample consisted of 35 participants (18 female, 16 male, 1 non-binary, M_age = 21.14 SD_age = 3.23, 5 left handed; see **Supplementary Table 1** for demographic information).

A power analysis showed that sample sizes of 25 and 30 participants would provide enough power to detect a moderate effect size in terms of standard difference scores (dz) of .51 and .46, respectively, in a one-tailed t-test (corresponding to fMRI contrasts between two conditions) with alpha = .05, uncorrected for multiple comparisons, and power = .80. Thus, our final sample size of 35 would provide enough power to detect a dz of .43.

### 2.2 Overview of Tasks

#### 2.2.1 Memory encoding and retrieval tasks

The memory paradigm used here (**Figure 1**) is one originally developed to study a cognitive phenomenon termed relational luring (Ichien et al., 2023). At encoding, participants view sequentially presented word pairs and decide whether the two words are related. At retrieval, they view a series of word pairs and are asked whether the two words had been paired together at encoding. Critically, all words are presented at encoding; thus, accurate performance requires recollection of the specific stimulus pairing. Word pairs presented at retrieval are either Intact (the same compound stimulus as at encoding) or Recombined (associated with different words). The Recombined trials are divisible into (a) trials involving Familiar relations, that is, ones for which participants had seen other exemplars at encoding wherein a word pair exemplified the same abstract relation (e.g., category-exemplar, such as mammal-dog), and (b) trials involving Unfamiliar relations – that is, ones for which participants had not seen other exemplars of the same abstract relation at encoding. The trials are divided evenly amongst the three types: Intact, Recombined-Familiar, and Recombined-Unfamiliar. Prior research on this paradigm (Ichien et al., 2023) indicates that participants are more likely to false alarm to recombined word pairs involving familiar than unfamiliar word pairs – a phenomenon referred to as relational luring (Popov et al., 2017).

**Figure 1:**
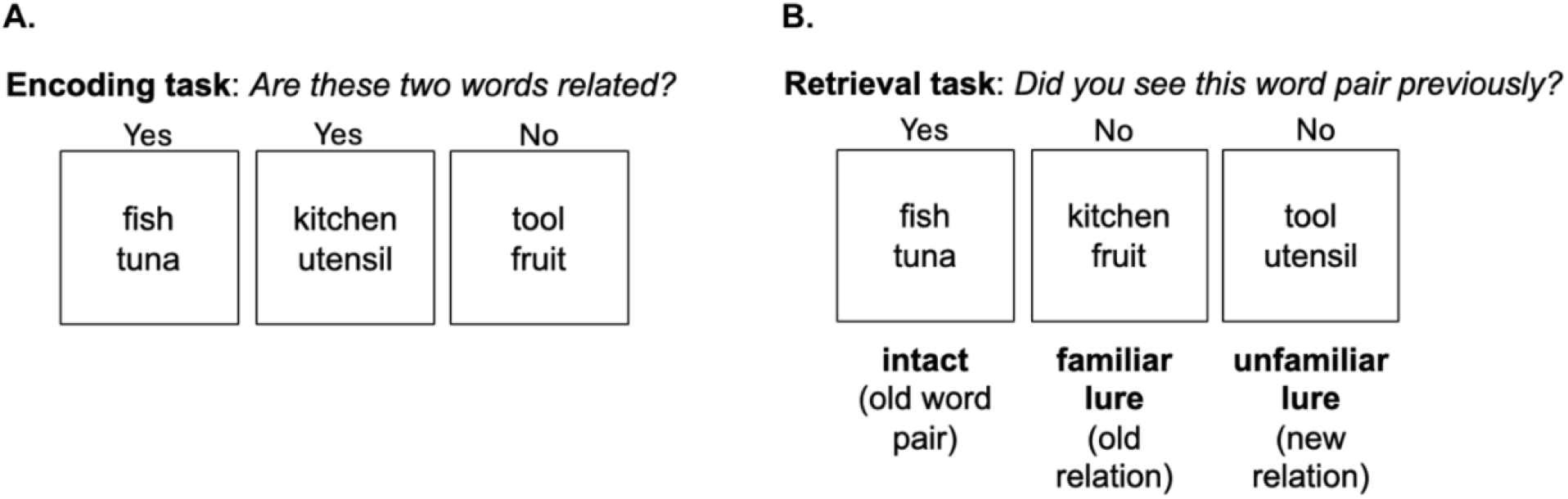
Overview of Memory encoding and retrieval tasks. **(A)** Encoding components of the Memory paradigm. **(B)** Retrieval components of the Memory paradigm. Questions presented in italics briefly summarize the instructions for each task. Participants performed the encoding task behaviorally first, followed by fMRI data collection of the Reasoning task (see Figure 2) and then the retrieval task (referred to in fMRI analyses as the Memory task).

In prior behavioral research (Ichien et al., 2023), we found that memory judgments were faster and more accurate for Intact than Recombined trials, consistent with reduced monitoring demands (Achim & Lepage, 2005). We also found that memory judgments were, for many participants, faster and more accurate for Recombined-Unfamiliar than Recombined-Familiar trials, consistent with the phenomenon of relational luring. This pattern of behavioral results is consistent with greater retrieval monitoring demands on Recombined than Intact trials, and-–for many participants—on Recombined-Familiar than Recombined-Unfamiliar trials.

Accordingly, given prior research implicating aLPFC in retrieval monitoring, we predicted that aLPFC activation during the retrieval task (referred to simply as the Memory task in fMRI analyses) would show corresponding patterns of univariate activation, with stronger activation for Recombined than Intact trials – and potentially also for Recombined-Familiar than Recombined-Unfamiliar trials. In the present study, the two Recombined conditions did not differ in response times (RTs) at the group level, suggesting a lack of consistent difference in monitoring between the two Recombined conditions. Since evidence of differential monitoring demands between conditions is critical for the present study, Recombined-Familiar and Recombined-Unfamiliar conditions were, as specified in the preregistration (<u>osf.io/7q5dr/overview</u>), grouped into one condition (Recombined trials) and compared with Intact pairs.

#### 2.2.2 Reasoning tasks

The Reasoning task was a variant of the relational matching paradigm, a simplified pictorial analogy task that reliably engages aLPFC (Christoff et al., 2003; Smith et al., 2007; Bunge et al., 2009; Wendelken et al., 2009). In this paradigm (**Figure 2A**), participants are asked to make similarity judgments about a pair of stimuli (1st-order relational judgments) or relations between two pairs of stimuli (2nd-order relational judgments). Each stimulus has two relevant features, such as Type (animals or vehicles) or Location (found in the water or on land). On 1st-order trials, participants are prompted to indicate whether items in a pair match on a specific feature. For example, the cue “Location” instructs them to indicate whether or not two items are found in the same location). They are asked to make 1st-order relational judgments for each of two pairs of stimuli. On 2nd-order trials, the cue “Match” instructs them to indicate whether items in the two pairs match *in the same way* – that is, according to the same feature (for example, if the pictures in the first pair are both found in the water and those in the second pair are both found on land, the answer would be ‘yes’). As in most prior studies, which all involved visually presented objects, both conditions are visually identical; thus, the comparison of 2nd- vs. 1st-order trials manipulates relational complexity. In cases where participants make two 1st-order judgments or one 2nd-order judgment, the response demands are larger in the former.

**Figure 2:**
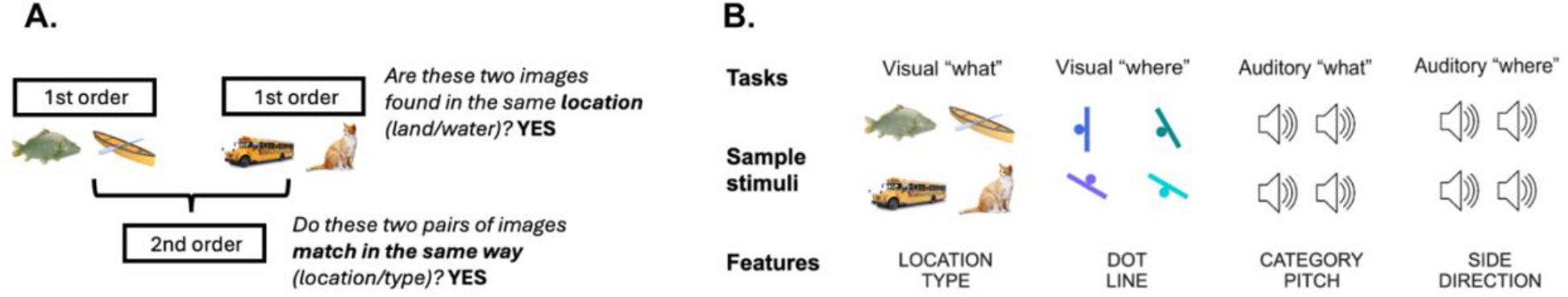
Overview of Reasoning tasks. **(A)** One of the four Reasoning tasks, involving visual-semantic content (Vis-What; see Supplement for analyses focused on this task). Questions presented in italics briefly summarize the instructions for each task. In each 1st-order trial, participants determined whether two stimuli matched along a cued dimension (eg., Location) (See Supplementary Figure 1 for a detailed account of the temporal structure). In each 2nd-order trial (cued as “Match”), participants determined whether two stimulus pairs matched on the same dimension. **(B)** Task-relevant features and example stimuli for each of the four Reasoning tasks: Vis-What (visual-semantic), Vis-Where (visual-spatial), Aud-What (auditory-featural), and Aud-Where (auditory-spatial).

The four relational reasoning tasks in the present study (**Figure 2B**) were developed as part of a larger study (OSF preregistration: <u>osf.io/fhjt2</u>; see also Vashel et al., in prep.). They all followed the same task structure and timing, with different types of stimuli and relevant stimulus dimensions: visual-semantic (“Vis-What”), visual-spatial (“Vis-Where”), auditory-featural (“Aud-What”), and auditory-spatial (“Aud-Where”).

### 2.3 Task Design and Stimuli

#### 2.3.1 Memory Encoding Task

##### 2.3.1.1 Task design

The encoding task was a 10-min task performed outside the scanner. It began with instructions on the screen, followed by a 5s fixation cross. Each word pair was presented for 2.5s, followed by a 1.5s fixation cross, with a total of 148 trials. Participants were given a break halfway through the task and were told to proceed when ready. Each half of the task was just under 5 minutes long.

##### 2.3.1.2 Stimuli

The stimulus set was constructed in such a way that 80% of word pairs were semantically related and instantiated one of three abstract semantic relations: category:exemplar (e.g., bird:robin), part:whole (e.g., toe:foot), and place:thing (e.g., store:groceries), while the other 20% of word pairs were not semantically related (e.g., mascara:spoon). A total of 148 word pairs were constructed out of 296 unique words. Word pairs were selected based on prevalence and concreteness norms (Brysbaert, Warriner, & Kuperman, 2014), as well as how well they instantiated the intended relations. Prevalence ratings were used to select words that are commonly known by most individuals. These norms provide a list of z-scores in which negative scores indicate that fewer than 50% of people know a word. Word pairs in our stimulus set only include words with a mean z-score of 2 or higher. Additionally, words included in the set had a mean concreteness score of at least 4 on a 1-5 scale, where a score of 5 indicates a concrete meaning (i.e., something that an individual can perceive directly with their senses) and a score of 1 indicates a highly abstract one.

To ensure that the word pairs instantiated the intended relation, we computed a Bayesian Analogy with Relational Transformations (BART) (Lu et al., 2012) typicality rating for each word pair, a process developed by Ichien et al. (2023). BART assumes that specific semantic relations between words are coded as distributed representations over a set of abstract relations. The BART model takes concatenated pairs of Word2vec vectors as the input and uses supervised learning with both positive and negative examples to acquire representations of individual semantic relations. After learning, the BART-based relational model calculates a relation vector consisting of the posterior probability that a word pair instantiates each of the learned relations (for details of the training procedure, see Ichien, Lu, & Holyoak, 2022). Only word pairs with a relational typicality rating (indicating how well the pair instantiates the abstract relation) of 0.5 or above were included in the encoding and retrieval tasks.

#### 2.3.2 Memory Retrieval Task

##### 2.3.2.1 Task design

The retrieval task was completed during one 9-minute scan, immediately preceded by the four reasoning tasks, and the encoding task before that. Participants were asked to identify whether or not they had seen that exact combination of words in the encoding task. The task began with instructions on the screen, followed by a 10s fixation cross. Each trial had a duration of 2.5s, followed by a jittered inter-trial interval that averaged 3s (range 2-6s), optimized for detection power using ‘easy-optimize-x’ (Spunt, 2016), a MATLAB-based design optimization tool. Responses were recorded within a 3-second window following the onset of the word pair. Participants completed 90 trials. The task ended with a 16s fixation cross.

##### 2.3.2.2 Stimuli

Participants completed an old/new recognition task in which they were presented with a sequence of 90 word pairs. Each word pair was constructed from individual words that participants had seen during their prior encoding task. Thus, each individual word was familiar to participants. A total of 90 word pairs were used for the retrieval task, with each word pair drawn from one of two broad types **(see Figure 1)**. The first type, Intact, consisted of word pairs that were shown during the encoding task, with the same number of word pairs instantiating each relation shown during the encoding task (i.e., category:exemplar, part:whole, and place:thing). The other type of word pair, Recombined, was constructed by recombining words that had appeared in the prior encoding task, so that individual words were now paired differently, generating novel word pairs distinct from those used in the encoding task.

The Recombined word pairs fell evenly into two subtypes: Recombined-Familiar and Recombined-Unfamiliar. Recombined-Familiar word pairs instantiated the same relations as the word pairs presented during the encoding task (e.g., two familiar words paired in a novel way to form a category:exemplar relation), with equal number of word pairs for each relation. Recombined-Unfamiliar word pairs instantiated a type of abstract relation (similarity) to which participants had not been exposed in the encoding phase. These word pairs were formed using words from the encoding set with overlapping salient attributes (e.g., bartender:cashier), and hence were relationally similar to one another, but not with respect to any of the three relations included in the encoding task. Among the 90 word pairs tested in the retrieval task, 30 pairs were Intact, 30 pairs were Recombined-Familiar, and 30 pairs were Recombined-Unfamiliar. As with the encoding task, each word pair had an average concreteness of 4 or above, based on concreteness norms from Brysbaert, Warriner, and Kuperman (2014). Additionally, both words in each word pair were presented in the same half of the encoding session, to prevent participants from being able to anchor their memory to the break in the middle.

In our prior study (Ichien et al., 2023), we noted that memory judgments for Recombined trials were more accurate when the first word in a pair shown at encoding (e.g., backyard:grill) was also presented first at retrieval (e.g., backyard:fruit). Thus, in constructing Recombined word pairs, we considered word position. All 30 Intact word pairs had both words in the same position at encoding and retrieval. For Recombined-Familiar and Recombined-Unfamiliar pairs, we ensured that roughly equal numbers of trials per condition (7 or 8 out of 30) had both words, the first word, the second word, or neither word in the same position relative to their position at encoding. In other words, the positions of both words were identical at encoding and retrieval for all Intact trials but for only around one quarter of Recombined trials; thus, mismatches in position could have increased retrieval monitoring demands on Recombined trials. Intact and Recombined trials were intermixed and presented in the same order for all participants in a given counterbalancing condition. Word pairs were pseudorandomly assigned across trials of a given type (Intact, Recombined).

#### 2.3.3 Reasoning tasks

##### 2.3.3.1 Task design

The reasoning paradigm was run as a blocked design, and completed across four runs of 9.57 minutes (9 min, 34 s) each. Each run included two of the four Reasoning tasks. The four combinations were Vis-What & Vis-Where, Vis-What & Aud-What, Vis-Where & Aud-Where, and Aud-What & Aud-Where.

For each trial, the cue indicating the dimension to focus on appeared at the onset. For example, in the case of the Vis-What task, either ‘Location’ or ‘Type’ appeared to cue the relevant dimension on 1st-order trials, and ‘Match’ appeared to cue 2nd-order trials. After 0.5s, the first stimulus image of each pair appeared for one second, followed by the second stimulus image of the pair 0.75s after the first image disappeared (See **Supplementary Figure 1** for temporal structure of a 1st-order trial). For 2nd-order trials, both pairs of images followed the same structure, with the two pairs presented sequentially. Although visual stimuli have been presented simultaneously in previous relational matching studies, this was not possible for auditory stimuli. Thus, both auditory and visual stimuli were presented sequentially to standardize timing across conditions and (in theory, at least) equate working memory demands. To orient participants to attend to the sound stimuli in the noisy scanner environment, a static playback icon was displayed on the screen concurrently with the presentation of each auditory stimulus. The icon carried no task-relevant information and was identical across trials.

Each 1st-order trial was 6s long and each 2nd-order trial was 12s long. Each trial was separated by 0.75s intertrial intervals (ITI) using a blank screen. Each 1st-order block consisted of four trials, and each 2nd-order block consisted of two trials. Each of the four runs included four 1st-order and four 2nd-order blocks for each of two tasks. Block types were interleaved, whereby each run started with a 1st-order block of a particular Reasoning task and then a 2nd-order, 1st-order, and 2nd-order block of the same task type, with blocks separated by a fixation cross, jittered between 8 and 10s. This set of four blocks of one task was followed by four blocks of a different task, with the same interleaved structure. The order of the two tasks within each run was counterbalanced across participants. Each scan started and ended with a 16s fixation cross.

Of the 32 1st-order trials of each task, there was an equal number of trials for each of the two features (eg., Location/Type for Vis-What task), and an equal number of trials with yes/no as the correct response. Of the 16 2nd-order trials for each task, 5 trials matched on one feature (eg., Location), 5 matched on the other feature (eg., Type), and 6 did not match on either feature.

##### 2.3.3.2 Stimuli

The stimuli and stimulus-generation procedures described in this section are also reported in Vashel et al. (in prep).

###### 2.3.3.2.1 Aud-What

The Aud-What condition required judgments based on pitch and instrument category while minimizing reliance on auditory spatial information. Stimuli consisted of recordings of musical instruments producing single notes. Instruments belonged to one of two categories—strings (violin, cello) and percussion (marimba, xylophone)—selected to maximize category discriminability, in the interest of standardizing task difficulty across stimulus types. Notes were drawn from four pitch classes (D, A♭, F, and B), each presented in both low (third octave) and high registers (fifth octave; fourth octave for B). Thus, the two task-relevant stimulus dimensions were instrument Category (string, percussion) and Pitch (low, high).

All auditory stimuli were obtained from publicly available sound libraries (Freesound and Pond5) and normalized to a duration of 1.0s. For percussion instruments, amplitude exhibited a natural decay across the stimulus interval. Twenty-eight unique stimuli were used in total, with individual stimuli repeated up to nine times across the experiment. Stimulus pairs were pseudorandomly generated, consisting of one low- and one high-register note, and filtered to preferentially include more dissonant pitch combinations. This constraint was introduced to increase perceptual discriminability and better equate task difficulty with the other stimulus types. Because pair selection was restricted by these requirements and we included only two instruments per category, up to six stimulus pairs were repeated; however, most pairs were unique, and no pair was repeated more than once.

###### 2.3.3.2.2 Aud-Where

The Aud-Where condition required judgments based on auditory spatial features while minimizing reliance on conceptual or semantic information. Stimuli simulated apparent sound motion either toward or away from the listener on the left or right side of auditory space. Accordingly, the two task-relevant dimensions were Direction (toward, away) and Side (left, right).

Stimuli were generated using the slab Python package (Schönwiesner et al., 2021). Binaural harmonic-complex tones were spatialized using KEMAR head-related transfer functions (HRTFs) interpolated at 0° elevation. Four azimuthal positions were defined on each side of auditory space (left: −110°, −80°, −70°, −40°; right: +40°, +70°, +80°, +110°). Each stimulus consisted of a sequence of four 250 ms tones presented at successive azimuthal positions, yielding a total duration of 1.0 s. Positions were ordered to create the percept of motion either toward or away from the midline. To reinforce perceived distance, tone amplitudes changed progressively across positions (multiplicative factors: 0.7, 0.8, 0.9, and 1.0), with lower amplitudes corresponding to more distant locations. Four fundamental frequencies (270, 280, 290, and 300 Hz) were used as a task-irrelevant distractor dimension. Frequencies were selected to fall within a comfortable audible range while minimizing interference from scanner noise. A total of sixteen unique stimuli were generated. Individual stimuli were repeated up to 14 times across the experiment, while all stimulus pairings remained unique.

###### 2.3.3.2.3 Vis-What

The Vis-What condition required judgments based on object identity and semantic attributes. Stimuli consisted of full-color photographs of familiar objects presented on a uniform white background. Objects varied along two task-relevant dimensions: Category (animal, vehicle) and Typical Location (land, water). Example animal stimuli included giraffe, moose, stingray, and sea turtle, whereas example vehicle stimuli included ferry boat, jet ski, fire truck, and motorcycle. Images were obtained from publicly available online repositories. A total of 57 unique stimuli were used, with individual stimuli repeated up to four times across the experiment. All stimulus pairings were unique.

###### 2.3.3.2.4 Vis-Where

The Vis-Where condition required judgments based on spatial features while minimizing reliance on conceptual or semantic information. Stimuli consisted of colored line segments presented at varying orientations, with a dot positioned on either the left or right side of the line. Accordingly, for relational matching, the two task-relevant dimensions were: Line (orientation of 30°, 60°, 90°, 120°, or 150°) and Dot (position on left or right side of the line). Stimuli were rendered in one of ten distinct colors, with color serving as a task-irrelevant distractor dimension to vary the stimuli sufficiently to make task difficulty more comparable to the other stimulus types. All visual stimuli were generated programmatically using a custom Python script. A total of 92 unique stimuli were used, with individual stimuli repeated up to four times. All stimulus pairings were unique.

### 2.4 Experimental Procedure

Prior to scheduling their session, participants were asked to watch an instructional video and complete a practice session online for the reasoning paradigm. The practice task was administered via Qualtrics and included two sample 2nd-order trials from the Vis-What task and two 2nd-order trials from the Vis-Where task. Participants were required to answer at least three of four practice questions correctly before being scheduled; they could repeat the practice task until they passed.

On the day of the session, participants first completed a short tutorial and seven practice trials for the encoding task. None of the words or relations used in the practice trials appeared in the experimental stimuli. Participants were instructed to respond even if the word pair disappeared from the screen, as responses were recorded until the next pair appeared. Participants then completed the encoding task outside the scanner. This was followed by a review of the Reasoning task instructions and an additional practice run consisting of 16 1st-order trials and 8 2nd-order trials spanning all four Reasoning tasks. Participants were required to reach a minimum performance threshold (i.e., four out of six correct for each task type and six out of eight correct on 2nd-order trials) before proceeding to scanning.

During MRI scanning, participants completed four runs of the relational reasoning paradigm, followed by one run of the retrieval task. Sounds were presented via Sensimetrics MRI-compatible insert earphones, and visual stimuli were projected onto a screen viewed through a mirror mounted on the head coil. Tasks were programmed and presented via PsychoPy (v2022.2.5). Participants responded using a button box held in their dominant hand. Right-handed participants used digits 2 and 3 to indicate “Yes” and “No,” respectively, whereas left-handed participants used digits 3 and 2 to indicate “Yes” and “No.”

### 2.5 Exclusion Criteria

Each paradigm had its own exclusion criteria, as outlined below, and participants were required to pass all criteria for both tasks to be included for analysis, as preregistered.

#### 2.5.1 Memory task exclusion criteria

Participants with a discriminability score (d’) below 1.68 on the encoding task were excluded, as we hoped to achieve a hit rate of 80% and a false alarm rate of 20%. Participants with a d’ score below zero on the Memory task were also excluded, as this indicated that the participant was not able to discriminate between Intact vs. Recombined word pairs. Participants were also excluded if they had more than 15% of TRs in the Memory task flagged as outlier volumes (>0.5 mm framewise displacement or >1.5 standardized DVARS).

#### 2.5.2 Reasoning task exclusion criteria

Behavioral exclusion criteria for the Reasoning task were applied at the participant, stimulus-type, and block levels. At the participant level, participants were required to contribute data from at least five of eight blocks for each task condition, and achieve at least 70% accuracy on 1st-order as well as 2nd-order trials. At the stimulus-type level, participants were required to achieve at least 62.5% accuracy for each of the four tasks, corresponding to performance above chance for the number of trials in each condition. At the block level, 1st-order blocks were retained if participants responded correctly on at least three of four trials, and 2nd-order blocks were retained if at least one of two responses was correct.

Motion-related exclusions were assessed at the block level. Motion outlier volumes (>0.5 mm framewise displacement or >1.5 standardized DVARS) were assigned to blocks based on their hemodynamic response function (HRF)-weighted contribution to the corresponding task-condition regressor. Specifically, a volume was associated with a block if it fell among the 12 volumes with the greatest contribution to that block. Blocks containing six or more motion outlier volumes (≥50% of associated volumes) were excluded. This threshold was selected based on empirical analyses indicating degradation of block-level beta estimates at higher levels of motion contamination.

### 2.6 ROI definitions

We focused on lateral subregions of area 10 (a10p and p10p), as well as adjacent regions a47r and p47r, which together comprise the aLPFC parcels of interest (Figure 3). We selected these parcels based on converging evidence from prior work implicating them in relational reasoning and higher-order cognitive control. Meta-analytic evidence from Neurosynth using the term “relational” highlights consistent engagement of peak coordinates that fall into a47r, p47r, and a10p. Moreover, a prior study (Assem et al., 2020) had identified a47r and p47r as regions preferentially engaged during relational reasoning, in contrast to “core” MDS regions, which are consistently activated across many cognitively demanding tasks. They described a47r and p47r as part of the MDS “penumbra”, a set of regions that show more task-selective recruitment and may support specialized computations depending on the cognitive demands. Furthermore, a47r and p47r have also been included in the definition of the frontal pole in some prior work (Baker et al., 2018).

**Figure 3.**
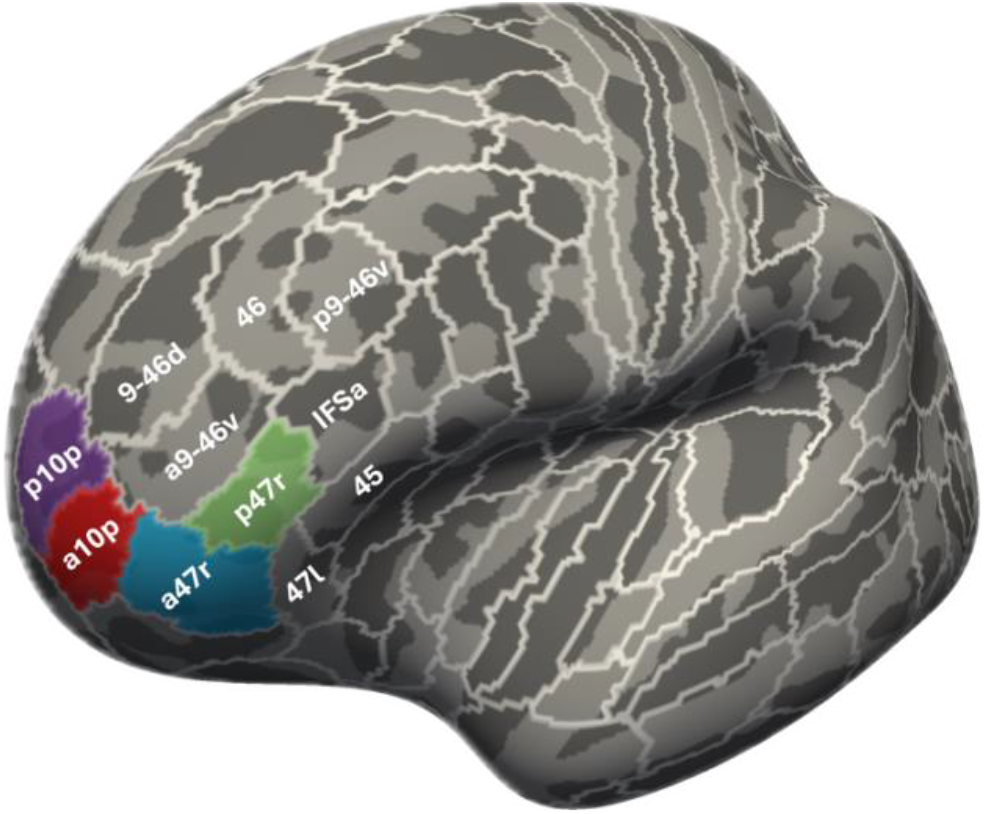
Left aLPFC parcels. Four parcels from the Glasser multimodal cortical parcellation (Glasser et al., 2016) spanning left aLPFC. Colors indicate parcels of interest selected for analysis.

We included three additional parcel definitions, all in the left hemisphere. First, given our expectations that functional activation patterns would not conform precisely to the parcel boundaries, and that the spatial distribution of activation would vary across individuals, we also defined the broader aLPFC region of interest comprising the union of a47r, p47r, a10p, and p10p. This allowed us to assess overlap at a broader spatial scale that is less sensitive to individual variability in functional topography. Second, we included primary visual cortex (V1) as a perceptually driven benchmark region that we expected would be highly sensitive to stimulus and task characteristics. Third, we included DLPFC parcel a9-46v as another benchmark; given that this general area has been implicated in domain-general cognitive control (Fedorenko et al., 2013) and that it has been designated as a “core” region of the MDS (Assem et al., 2020; Duncan et al., 2020), we expected it to be relatively insensitive to stimulus and task characteristics. .

### 2.7 MRI Data Acquisition

Whole-brain imaging was conducted on a Siemens 3.0T Prisma MRI scanner at the Henry H. Wheeler, Jr. Brain Imaging Center at University of California, Berkeley. Using a 32-channel head coil, functional images were acquired using an SMS (multiband) EPI sequence (factor = 3; TR = 2000 ms; TE = 34 ms; voxel size = 2 mm isotropic; flip angle = 66°; matrix dimensions = 104 × 104 × 75 voxels). Functional data were collected across four runs of 287 volumes (9.56 minutes) each for the relational reasoning paradigm, followed by one run of 270 volumes (9 minutes) for the Memory task. Four initial volumes from each run were discarded to allow for scanner stabilization prior to task onset. High-resolution structural volumes were collected using a 3D, T1w MPRAGE sequence at 0.9-mm isotropic resolution (TR = 2300 ms, TE = 2.32 ms, flip angle = 8°; matrix dimensions = 256 × 256 × 192 voxels).

### 2.8 fMRI Preprocessing

Raw DICOM images were converted to NIfTI format using dcm2niix.

Results included in this manuscript come from preprocessing performed using *fMRIPrep* 24.1.1 (Estoban et al., 2018; Estoban et al., 2019; RRID:SCR_016216), which is based on *Nipype* 1.8.6 (Gorgolewski et al., 2011; Gorgolewski et al., 2018; RRID:SCR_002502).

Preprocessing of B_0_ inhomogeneity mappings: A total of 5 fieldmaps were found available within the input BIDS structure for this particular subject. A *B0*-nonuniformity map (or *fieldmap*) was estimated based on two (or more) echo-planar imaging (EPI) references with topup (Andersson et al., 2003; FSL None).

Anatomical data preprocessing: A total of 1 T1-weighted (T1w) images were found within the input BIDS dataset. The T1w image was corrected for intensity non-uniformity (INU) with *N4BiasFieldCorrection* (Tustison et al., 2010), distributed with ANTs 2.5.3 [Avants et al., 2008, RRID:SCR_004757], and used as T1w-reference throughout the workflow. The T1w-reference was then skull-stripped with a *Nipype* implementation of the antsBrainExtraction.sh workflow (from ANTs), using OASIS30ANTs as target template. Brain tissue segmentation of cerebrospinal fluid (CSF), white-matter (WM) and gray-matter (GM) was performed on the brain-extracted T1w using ‘fast’ [FSL (version unknown), RRID:SCR_002823, Zhang et al., 2001]. Brain surfaces were reconstructed using ‘recon-all’ [FreeSurfer 7.3.2, RRID:SCR_001847, Dale et al., 1999], and the brain mask estimated previously was refined with a custom variation of the method to reconcile ANTs-derived and FreeSurfer-derived segmentations of the cortical gray-matter of Mindboggle [RRID:SCR_002438, Klein et al., 2017]. Volume-based spatial normalization to one standard space (MNI152NLin2009cAsym) was performed through nonlinear registration with ‘antsRegistration’ (ANTs 2.5.3), using brain-extracted versions of both T1w reference and the T1w template. The following template was were selected for spatial normalization and accessed with *TemplateFlow* [24.2.0, Ciric et al., 2022]: *ICBM 152 Nonlinear Asymmetrical template version 2009c* [Fonov et al., 2011, RRID:SCR_008796; TemplateFlow ID: MNI152NLin2009cAsym].

Functional data preprocessing: For each of the 5 BOLD runs found per subject (across all tasks and sessions), the following preprocessing was performed. First, a reference volume was generated, using a custom methodology of *fMRIPrep*, for use in head motion correction. Head-motion parameters with respect to the BOLD reference (transformation matrices, and six corresponding rotation and translation parameters) are estimated before any spatiotemporal filtering using ‘mcflirt’ [FSL, Jenkinson et al., 2002]. The estimated *fieldmap* was then aligned with rigid-registration to the target EPI (echo-planar imaging) reference run. The field coefficients were mapped on to the reference EPI using the transform. The BOLD reference was then co-registered to the T1w reference using ‘bbregister’ (FreeSurfer) which implements boundary-based registration [Greve & Fischl, 2009]. Co-registration was configured with six degrees of freedom. Several confounding time-series were calculated based on the *preprocessed BOLD*: framewise displacement (FD), DVARS and three region-wise global signals. FD was computed using two formulations following Power (absolute sum of relative motions, Power et al., 2014) and Jenkinson (relative root mean square displacement between affines, Jenkinson et al., 2002). FD and DVARS are calculated for each functional run, both using their implementations in *Nipype* [following the definitions by Power et al. (2014)]. The three global signals are extracted within the CSF, the WM, and the whole-brain masks. Additionally, a set of physiological regressors were extracted to allow for component-based noise correction [*CompCor*, Behzadi et al., 2007]. Principal components are estimated after high-pass filtering the *preprocessed BOLD* time-series (using a discrete cosine filter with 128s cut-off) for the two *CompCor* variants: temporal (tCompCor) and anatomical (aCompCor). tCompCor components are then calculated from the top 2% variable voxels within the brain mask. For aCompCor, three probabilistic masks (CSF, WM and combined CSF+WM) are generated in anatomical space. The implementation differs from that of Behzadi et al. (2007) in that instead of eroding the masks by 2 pixels on BOLD space, a mask of pixels that likely contain a volume fraction of GM is subtracted from the aCompCor masks. This mask is obtained by dilating a GM mask extracted from the FreeSurfer’s *aseg* segmentation, and it ensures components are not extracted from voxels containing a minimal fraction of GM. Finally, these masks are resampled into BOLD space and binarized by thresholding at 0.99 (as in the original implementation). Components are also calculated separately within the WM and CSF masks. For each CompCor decomposition, the *k* components with the largest singular values are retained, such that the retained components’ time series are sufficient to explain 50 percent of variance across the nuisance mask (CSF, WM, combined, or temporal). The remaining components are dropped from consideration. The head-motion estimates calculated in the correction step were also placed within the corresponding confounds file. The confound time series derived from head motion estimates and global signals were expanded with the inclusion of temporal derivatives and quadratic terms for each [Satterthwaite et al., 2013]. Frames that exceeded a threshold of 0.5 mm FD or 1.5 standardized DVARS were annotated as motion outliers. Additional nuisance timeseries are calculated by means of principal components analysis of the signal found within a thin band (*crown*) of voxels around the edge of the brain, as proposed by [Patriat et al., 2017]. The BOLD time-series were resampled onto the following surfaces (FreeSurfer reconstruction nomenclature): *fsaverage*, *fsnative*. All resamplings can be performed with a single interpolation step by composing all the pertinent transformations (i.e. head-motion transform matrices, susceptibility distortion correction when available, and co-registrations to anatomical and output spaces). Gridded (volumetric) resamplings were performed using nitransforms, configured with cubic B-spline interpolation. Non-gridded (surface) resamplings were performed using mri_vol2surf (FreeSurfer).

Many internal operations of *fMRIPrep* use *Nilearn* 0.10.4 [Abraham et al., 2014, RRID:SCR_001362], mostly within the functional processing workflow. For more details of the pipeline, see the section corresponding to workflows in *fMRIPrep*’s documentation (https://fmriprep.readthedocs.io/en/latest/workflows.html).

The above boilerplate text was automatically generated by fMRIPrep with the express intention that users should copy and paste this text into their manuscripts unchanged. It is released under the CC0 license.

### 2.9 General linear modeling of fMRI data

Functional MRI data were analyzed using first-level general linear models (GLMs) implemented in Nilearn (Abraham et al., 2014). For each participant, the first four TRs from each functional run of each task were removed to allow for stabilization of the scanner. The four runs from the Reasoning task were concatenated for analysis.

A design matrix for each task was constructed for each participant using Nilearn. For the Memory task, we used four task regressors of interest, corresponding to the two task conditions (Intact, Recombined), crossed with trial-level correctness (Correct, Incorrect), creating a trial-level GLM. For the Reasoning task, we used eight task regressors of interest, corresponding to the eight task conditions defined by relational complexity (1st-order, 2nd-order) crossed with stimulus set (Aud-What, Aud-Where, Vis-What, Vis-Where), creating a block-level GLM. Task regressors were convolved with the canonical SPM HRF. Because fMRIPrep performs slice-time correction relative to the middle of each TR, accounting for simultaneous multi-slice (multiband) acquisition, event onsets were shifted by one-half TR prior to construction of the design matrix to maintain temporal alignment between task regressors and the preprocessed BOLD signal. In the Reasoning task, blocks that did not meet predefined behavioral performance or motion quality criteria were modeled separately as excluded blocks.

Nuisance regressors included six rigid-body motion parameters, the first five anatomical CompCor components (Behzadi et al., 2007), discrete cosine drift terms, run-specific intercepts, and binary regressors for motion-outlier volumes. Motion outliers were defined by fMRIPrep as volumes exceeding 0.5 mm framewise displacement or 1.5 standardized DVARS. In the Memory task, temporal derivatives of each task regressor were included to account for variability in the timing of the hemodynamic response. For the Reasoning task, to account for variation in BOLD response related to task difficulty or time-on-task effects, an additional response-time regressor was included, defined as the average response time for each block and convolved with the canonical HRF (Grinband et al., 2008).

First-level model estimation was performed using vertex-wise ordinary least squares GLMs implemented in Nilearn. All analyses were conducted using surface-based representations. Surface-based analyses were used because they provide improved anatomical alignment and statistical sensitivity relative to volume-based approaches (Spencer et al., 2022).

Individual-level overlap analyses were conducted in fsnative space, whereas group-level univariate analyses were conducted in fsaverage space to facilitate vertex-wise comparisons across participants. Participant-specific Glasser parcel labels were generated by projecting the HCP-MMP1.0 atlas from fsaverage space to each participant’s native cortical surface using the surface registration mappings generated during fMRIPrep processing.

### 2.10 fMRI Analysis

#### 2.10.1 Measurement of Task Overlap

We computed Memory and Reasoning task contrasts for each participant. The memory contrast consisted of correct Recombined > correct Intact trials, and the reasoning contrast of 2nd-order > 1st-order reasoning blocks. We then quantified task overlap in several ways. First, we conducted group-level whole-brain contrasts to identify regions engaged in both tasks or only one, and determined which prefrontal parcels fell into one of these categories. Second, we conducted ROI analyses at the individual level to test whether clusters engaged during the Reasoning task by a given individual also showed sensitivity to the Memory task manipulation. Third, to assess overlap between tasks at a finer spatial scale, we quantified the degree of overlap in patterns of activation across the most active vertices within each aLPFC parcel. Fourth, we conducted exploratory individual differences analyses testing whether degree of neural overlap was correlated with memory performance across individuals.

For our primary analyses, we had preregistered that we would measure overlap between the Memory task and a composite of all four Reasoning tasks if the latter showed strongly overlapping activation to one another; as this was the case, we report these analyses here. We additionally examined overlap between the Memory task and one of the four relational matching tasks––the Vis-What task––to roughly equate the amount of data included across tasks (**Supplementary Figure 9**). As noted in the preregistration, we selected the Vis-What task because its content was most comparable to the Memory task: while the former involved pictures and the latter involved words, both required consideration of semantic relations among visually presented stimuli.

##### 2.10.1.1 Group-level analyses

To visualize our contrasts of interest at the group level, we computed whole-brain group-level *t*-tests for the memory and reasoning contrasts separately, setting a significance threshold of *t* ≥ 2.5. Then, to examine shared and distinct activation across tasks, we computed conjunction and disjunction maps. Our conjunction analyses identified regions significantly active in both contrasts (*t* ≥ 2.5), whereas our disjunction maps identified regions showing task-specific activation. For each disjunction analysis, we imposed a threshold of *t* ≥ 2.5 for the task of interest and |*t*| < 1 for the other task to indicate insensitivity to that task manipulation in either direction (i.e., ensuring that 2nd-order is neither significantly higher or lower than 1st-order; Recombined is neither significantly higher or lower than Intact). We noted which aLPFC parcels had significant vertices for these analyses.

##### 2.10.1.2 Individual-level analyses

To complement group-level findings, we conducted analyses at the individual level in the left hemisphere (see Supplement for right hemisphere results). These analyses focused on (i) testing whether individually defined clusters engaged during reasoning were also sensitive to memory retrieval demands and linked to memory performance across individuals, and (ii) quantifying overlap in the most active vertices across tasks.

###### 2.10.1.2.1 Functionally defined clusters

We first identified clusters of activation from the Reasoning task contrast, defined *a priori* as sets of at least 50 contiguous vertices exceeding a threshold of *z* ≥ 2. For each participant, we quantified the number of clusters within each parcel, as well as within a broader aLPFC parcel comprising all four parcels (a47r, p47r, a10p, p10p). For clusters spanning parcel boundaries, only vertices within the aLPFC parcels were retained, and the cluster was assigned to the parcel containing the greatest number of vertices. We then identified the two parcels that had the highest proportion of participants exhibiting clusters of at least 50 vertices (74% and 49%, as opposed to 37% and 34%). For these two parcels, we extracted beta estimates from the Memory task (Recombined vs. Intact contrast) within each participant’s individually defined clusters. To summarize activation at the participant level, we computed weighted beta estimates, yielding a single value per participant.

###### 2.10.1.2.2 Individual-level patterns of active vertices

For each participant and each paradigm, we identified the top 5% of most active vertices within each parcel. We then computed the Dice coefficient, a measure of spatial overlap ranging from 0 (no overlap) to 1 (perfect overlap), to quantify the degree of correspondence between tasks.

To determine whether observed overlap exceeded chance levels, we generated participant-specific null distributions. For each participant and parcel, we permuted the vertex labels of the Reasoning task activation map 1,000 times and recomputed Dice coefficients for each permutation. Because parcels varied in size across participants, null distributions were computed separately for each participant and parcel.

We then quantified evidence for above-chance overlap using Bayes factors (BF10), following guidelines described by Kelter (2020). Specifically, we computed Bayes factors based on the difference between each participant’s observed Dice coefficient and the mean of their null distribution. Bayes factors were computed for each of the four aLPFC parcels, the broader aLPFC ROI, and the two comparison regions: V1 and a9-46v. These comparisons provided benchmarks for interpreting the specificity of overlap observed in aLPFC.

#### 2.10.2 Exploratory correlations with memory performance

As an exploratory analysis to assess whether activation in these clusters identified from the Reasoning task was behaviorally meaningful in the Memory task, we examined correlations between cluster-level activation and memory performance, quantified using *d′* (d-prime) to account for response bias arising from unequal proportions of trial types (twice as many trials on which the correct answer was “no” (Recombined word pairs) relative to “yes” (Intact word pairs). We additionally examined correlations between Dice coefficients in each parcel and memory performance (*d’*) to assess whether the degree of vertex-level overlap between the two tasks was behaviorally meaningful.

#### 2.10.3 Projection of published coordinates to the cortical surface

To assess the correspondence between previously reported volumetric coordinates and the surface-based Glasser parcellation, we projected spherical ROIs centered on the published MNI coordinates from MNI152 volumetric space onto the FreeSurfer fsaverage surface using *mri_vol2surf* and the MNI152-to-fsaverage registration. The resulting surface projections were thresholded and clustered to identify contiguous cortical regions, which were then compared with the Glasser multimodal parcellation to determine the parcels encompassed by each projected ROI.

## 3 RESULTS

### 3.1 Behavioral Results

#### 3.1.1 Memory Task

Overall accuracy was high at encoding and more variable at retrieval. At encoding, accuracy ranged from 0.82 to 1.0 (MAcc = 0.95, SDAcc = 0.04). Discriminability (*d’)* ranged from 2.20 to 4.22, with a mean of 3.43. At retrieval, accuracy was higher for Intact pairs (MAcc = 0.82, SDAcc = 0.11) compared to Recombined pairs (MAcc = 0.65, SDAcc = 0.21), *t*(34) = 3.83, *p* < .001, *dz* = 0.65. Discriminability (*d’*) ranged from 0.13 to 3.03, with a mean of 1.45. Excluding participants with d’ < 1 did not change the pattern of results, so all participants were included in subsequent analyses. As predicted, RTs were longer for Recombined pairs (MRT = 1.61 s, SDRT = 0.27) than Intact pairs (MRT = 1.30 s, SDRT = 0.28), t(34) = 9.88, *p* < .001, *dz* = 1.67. Having operationalized longer RTs at retrieval as reflecting higher retrieval monitoring demands, we considered this evidence of heightened monitoring on Recombined than Intact trials.

Among Recombined trials, accuracy was higher for Recombined-Unfamiliar pairs than Recombined-Familiar pairs, *t*(34) = 2.56, *p* < 0.05, *dz* = 0.43, but RTs did not differ significantly between these conditions, *t*(34) = -1.58, *p* = 0.123, *dz* = -0.27. Because of the absence of an effect on RTs, we did not have evidence of greater monitoring demands on Recombined-Familiar trials; thus, as planned, we combined Recombined-Familiar and Recombined-Unfamiliar trials in subsequent analyses. An exploratory analysis confirmed minimal LPFC activation (with no aLPFC activation) for a direct contrast between these conditions.

#### 3.1.2 Reasoning Task

Participants performed at near-ceiling accuracy on both tasks (1st-order trials: MAcc = 0.96, SDAcc = 0.03, range = 0.91-1.00; 2nd-order trials: MAcc = 0.95, SDAcc = 0.06, range = 0.73-1.00). A paired-samples t-test indicated no significant difference in accuracy between conditions, *t*(34) = 1.01, p = 0.32, *dz* = 0.17. Response times (RTs), measured from the onset of the final stimulus in each trial (second stimulus for 1st-order, fourth stimulus for 2nd-order) until the end of the ITI, differed between levels of relational complexity. Mean RTs were faster for 2nd-order trials (MRT = 1.05 s, SDRT = 0.25, range = 0.68-1.74) than 1st-order trials (MRT = 1.18 s, SDRT = 0.21, range = 0.76-1.52), with a significant difference, *t*(34) = 6.20, *p* < .001, *dz* = 1.05. In sum, the two conditions did not differ on accuracy, and RTs were in fact faster on the condition involving higher relational complexity, highlighting the fact that the conditions are cognitively demanding in different ways in this variant of the relational matching paradigm (see Kelly et al., 2026).

### 3.2 fMRI Results

#### 3.2.1 Group-level overlap in activation

To characterize task activation at the group level, we computed whole-brain group contrasts (**Figure 4**) for the Reasoning task (2nd > 1st-order blocks) and Memory task (correct Recombined > correct Intact trials). We computed two-tailed t-tests with a threshold of |*t*| ≥ 2.5, as opposed to the preregistered one-tailed t-test with a threshold of *p* < .01.

**Figure 4.**
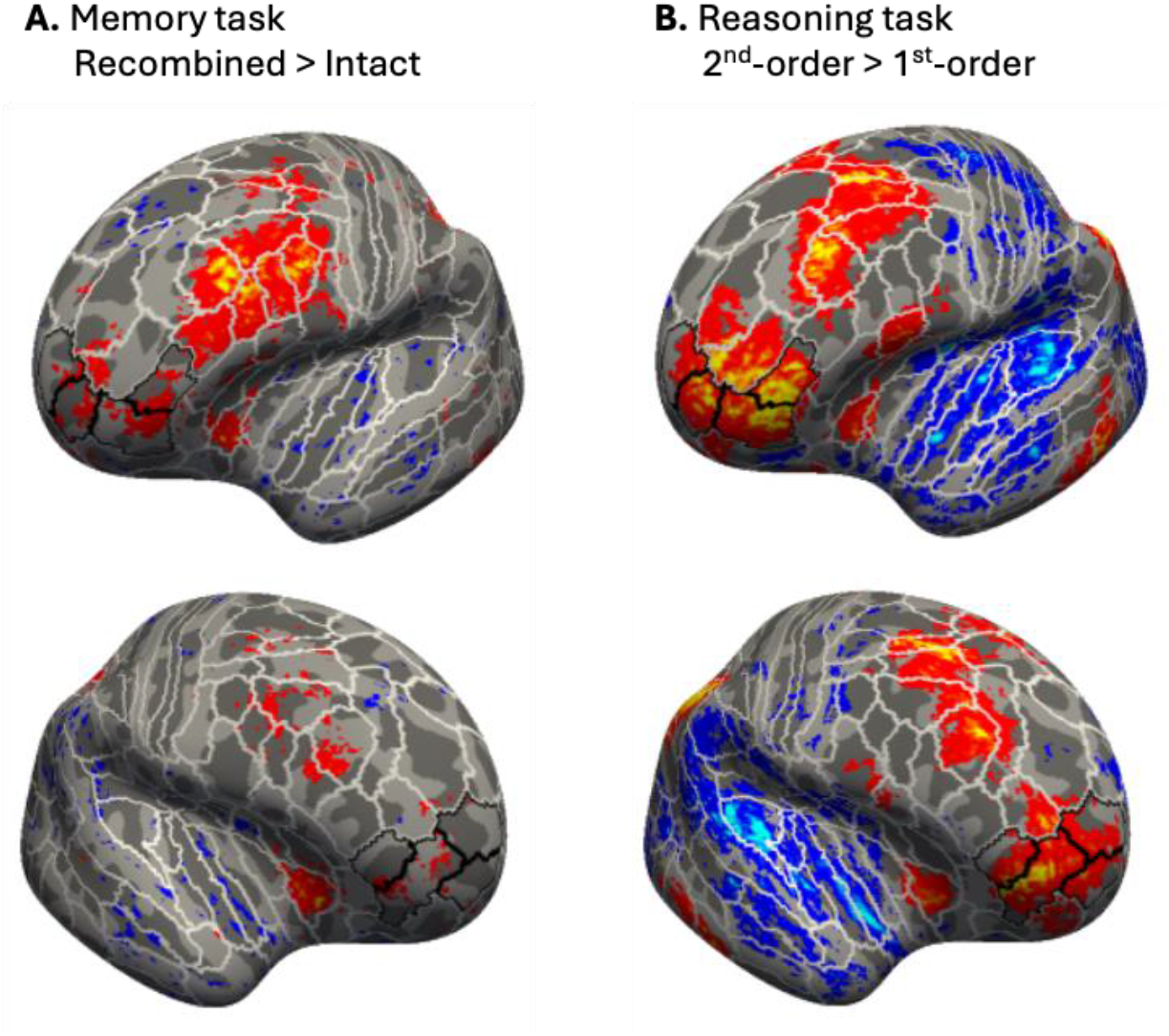
Whole-brain group-level activation contrasts during the memory and reasoning paradigms. aLPFC parcels are outlined in black. **(A)** Recombined > Intact contrast and **(B)** 2nd-order > 1st-order contrast, showing left and right hemispheres. Activation contrasts were computed with a threshold of |t| ≥ 2.5.

The Reasoning task manipulation engaged a broad network within left PFC, including all four aLPFC parcels of interest (a47r, p47r, a10p, p10p), as well as DLPFC parcels a9-46v, p9-46v, and 9-46d. In the right hemisphere, activation was observed across the same parcels; indeed, right-hemisphere activation was more robust than has been reported in most studies (see Neurosynth meta-analysis for the term “reasoning” or “relational”; Yarkoni et al., 2011), perhaps because of the large amount of data collected here (∼40 minutes).

The Memory task manipulation engaged all four parcels of interest in left aLPFC, along with dorsolateral parcels a9-46v and p9-46v, and a ventrolateral parcel: anterior inferior frontal sulcus (IFSa). Activation was not as robust as for the Reasoning task, which was to be expected given that the combination of all four Reasoning tasks included more than four times as much data. That said, even when the amount of data was comparable (see Section 3.2.3; **Supplementary Figure 8**), activation was still more robust for the Reasoning manipulation; this may be because the Memory manipulation was more subtle, and that the task was event-related design as opposed to a blocked design. In the right hemisphere, activation was more limited and was observed largely in aLPFC parcel a47r and DLPFC parcel p9-46v.

Next, to identify overlapping regions at the group level, we performed a conjunction analysis of the Reasoning and Memory task contrasts (**Figure 5A**). This analysis revealed overlapping activation in left PFC within all four aLPFC parcels of interest (a47r, p47r, a10p, p10p), extending into DLPFC parcels a9-46v and p9-46v. In the right hemisphere, overlap was observed in aLPFC parcels a47r, p47r, and a10p, as well as DLPFC parcels a9-46v, p9-46v, and 9-46d. Thus, there was overlapping group-level activation across tasks in all areas of interest and beyond.

**Figure 5.**
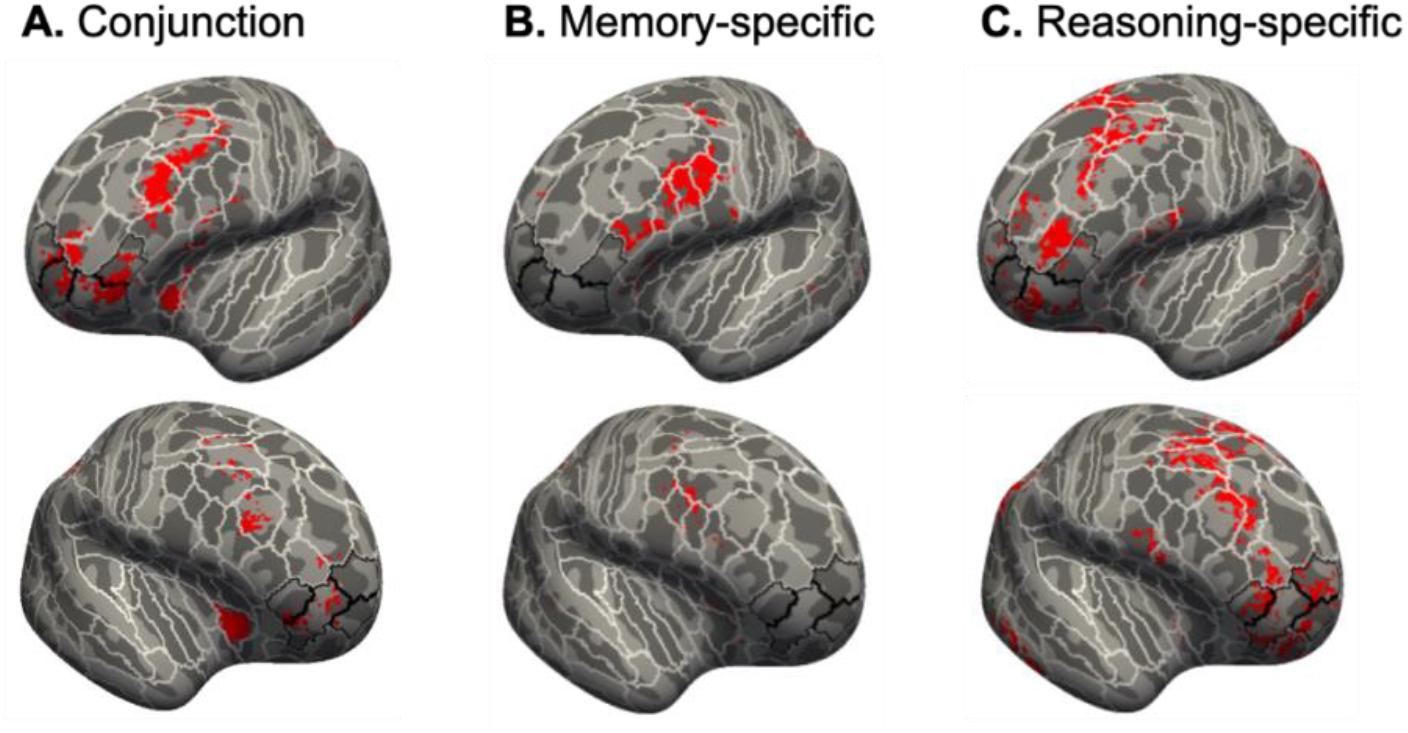
Cortical areas showing overlapping and non-overlapping activation across memory and reasoning paradigms. aLPFC parcels are outlined in black. **(A)** Conjunction showing overlap of activation between memory and reasoning paradigms in the left and right hemispheres, computed as the intersection of both activation contrast maps, with a threshold of t ≥ 2.5. **(B)** Disjunction showing memory-specific activation. **(C)** Disjunction showing reasoning-specific activation. Disjunction maps were computed as the intersection of task-specific activation maps, with a positive threshold of t ≥ 2.5 in one task, to indicate significant activation, and a threshold of |t| < 1 for the other task, to indicate lack of significant activation.

In addition to identifying areas of overlap, we performed group-level disjunction analyses identifying regions that were engaged in one task but not the other. The reasoning-specific analysis (**Figure 5C**) revealed activation in sparse clusters within left aLPFC, alongside activations in DLPFC: a prominent cluster in left a9-46v and more posterior regions spanning 9-46d, 46, and p9-46v. A similar pattern was observed in the right hemisphere. In contrast, the memory-selective disjunction analysis (**Figure 5B**) revealed no activation in left aLPFC; this null effect was unlikely due to insufficient sensitivity of the Memory task, given that a large left LPFC cluster was identified in the left hemisphere that spanned multiple parcels, including VLPFC parcel left IFSa. Moreover, the lack of aLPFC activation in the Memory disjunction does not stem from the imbalance in amount of data included for the two tasks, as this was also observed when restricting the Reasoning contrast to a single task (**Supplementary Figure 9**).

#### 3.2.2 Individual-level analyses

To move beyond these group-level results, we next conducted analyses at the individual level – both via ROI analyses and quantification of patterns of overlap. First, we tested whether individually identified clusters of activation from the reasoning contrast were also sensitive to memory retrieval demands. Second, we quantified overlap in the most active vertices across tasks. As exploratory analyses, we also tested whether the reasoning-related clusters of activation and the degree of vertex overlap were linked to memory performance. While we focused on the left hemisphere, for which we had clearer predictions, we also report results for the right hemisphere (**Supplementary Figures 4-7**).

##### 3.2.2.1 Cluster-based ROI analyses

We first identified parcels for which a substantial proportion of participants exhibited clusters of activation for the Reasoning task manipulation (2nd > 1st-order blocks), defined as ≥ 50 contiguous vertices with *z* > 2 (**Supplementary Figure 2**). While 80% of participants exhibited at least one cluster of this size within the broader left aLPFC region, this proportion was lower when considering individual parcels: 60% in a47r, 48.6% in p47r, 31.4% in a10p, and 25.7% in p10p, because of individual variability in terms of which parcels the clusters were observed in – that is, variability in functional organization that is not captured by this group-level parcellation scheme. Notably, only one participant showed significant clusters of task activation (2nd-order blocks > baseline) at this statistical threshold in a10p or p10p (**Supplementary Figure 3**) even though they showed differential activation for 2nd-order > 1st-order trials, indicating limited task-positive engagement relative to the resting baseline in these clusters. Cluster prevalence was lower in the right hemisphere, but showed a similar pattern across parcels (Section 3.2.2.4; see **Supplementary Figure 2**). Given that we sought to identify regions with relatively more consistent activation during reasoning, subsequent cluster-based analyses focused on left a47r and p47r, as well as the broader left aLPFC region.

To test whether reasoning-related clusters within these parcel definitions were sensitive to memory demands, we conducted paired t-tests comparing mean beta values in these clusters for correct Recombined vs. Intact trials (**Figure 6A**). Stronger activation for Recombined trials was observed in all three parcel definitions (t-values 3.0 - 4.3; all p < .01). All effects remained significant after correcting for multiple comparisons. To determine whether these clusters (identified across the full reasoning dataset) were sensitive to stimulus domain, and in particular whether any particular task could be driving the activation, we also derived mean beta values from each individual task and conducted a 2 (relation level: 1st vs. 2nd-order) × 4 (task) repeated-measures ANOVA (shown for the full aLPFC parcel in **Figure 6B**). There was—as expected based on how the clusters were derived—a robust main effect of relation level; to the point, however, we found that activation parameters were comparable across tasks. Likewise, aLPFC sensitivity was more robust for the Reasoning manipulation than the Memory manipulation, which was also to be expected in part based on how the clusters were derived. Importantly, the fact that the clusters derived from the Reasoning manipulation were sensitive to the Memory manipulation cannot be explained by similarity in stimulus processing demands, given that the tasks spanned several sensory modalities and stimulus domains. In summary, ROIs individually derived from reasoning-related clusters (of at least 50 contiguous vertices at Z > 2) were most consistently observed in a47r and p47r; these clusters were sensitive to the Memory manipulation.

**Figure 6.**
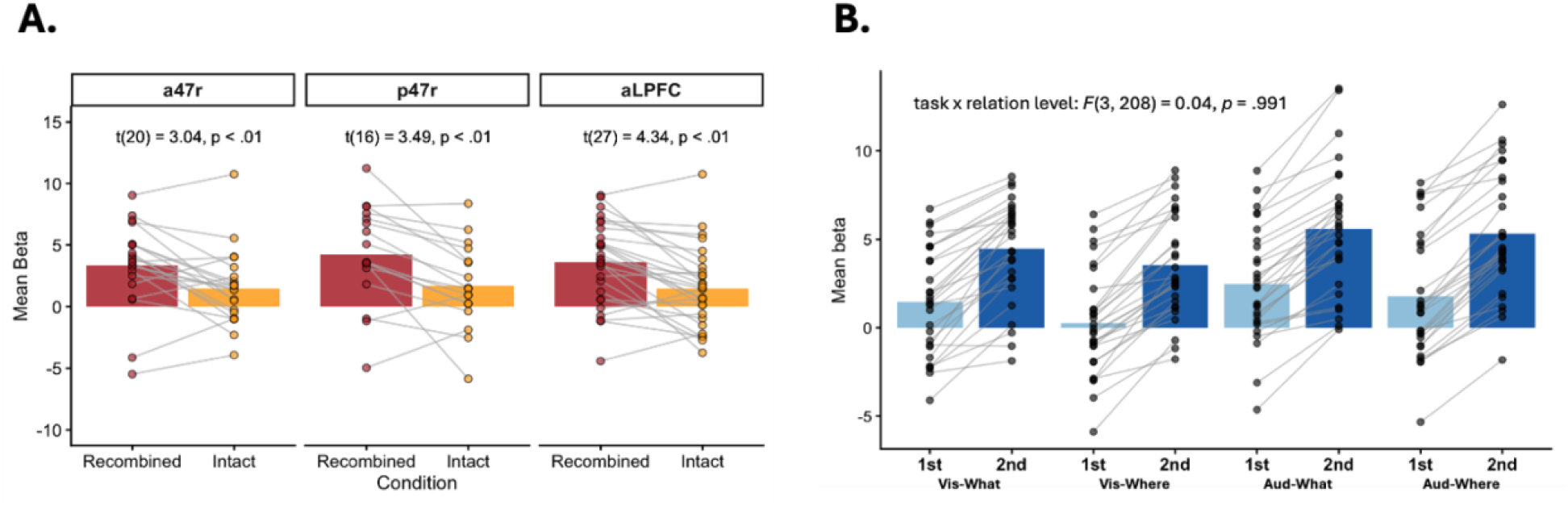
Sensitivity to memory retrieval demands and relational complexity in reasoning-related clusters in left aLPFC. **(A)** Activation in reasoning-related clusters during memory retrieval. Cluster-weighted mean beta values are shown for Correct Recombined and Correct Intact trials. Points represent individual participants, and lines connect participants across conditions. Paired t-test results are shown for each parcel definition. Note that fewer participants had significant clusters in the individual parcels than in aLPFC as a whole. **(B)** Sensitivity to relational complexity in reasoning-related clusters across all four Reasoning tasks in aLPFC. Clusters were defined using data from the entire reasoning paradigm and therefore are inherently biased; however, this analysis verified that observed effects were not driven solely by the Vis-What task. Points represent cluster-weighted mean beta values for individual participants, and bars indicate the mean across participants.

##### 3.2.2.2 Overlap in active vertices

To quantify overlap at a more fine-grained level, we identified the top 5% of active vertices in the left hemisphere for each contrast (2nd > 1st-order; Recombined > Intact) within each parcel and participant, ensuring that these vertices all showed above-zero beta weights for these contrasts. Here, we examined all four individual aLPFC parcels because—unlike for the cluster analysis— spatially contiguous activation across vertices was not a prerequisite for these vertex-level analyses. Mean Dice coefficients across aLPFC parcels ranged from 0.12 in p10p to 0.24 in p47r (0.17 to 0.24 before removing outlier participants, separately for each parcel); in other words, on average, 12 to 24% of the very most active vertices within a parcel overlapped between tasks. The overlap for all aLPFC parcels was numerically greater than for either benchmark regions (V1 parcel: mean Dice = 0.05; a9-46v parcel: mean Dice = 0.17).

Notably, there was substantial variability across participants. For example, Dice coefficients within p47r for the top 5% of vertices ranged from 0.0 to 0.79 (M = 0.24, SD = 0.04) – i.e., for some participants, none of the most active vertices overlapped in this parcel, whereas for others, 79% did. Inter-individual variability was also high in the broader aLPFC region, with Dice coefficients varying from 0 to 59% for the top 5% of vertices.

When expanding to consider larger proportions of top vertices, the degree of overlap increased. Mean Dice coefficients rose when including the 10% or 30% most active vertices, reaching a maximum of 41% average overlap in p47r at the 30% threshold. However, inter-individual variability decreased only slightly when expanding the number of top vertices included, and the overall pattern across parcels obtained at 30% largely mirrored those obtained at 5%.

Given the substantial individual variability, we sought to test whether the degree of overlap was greater than expected by chance. To assess whether the observed overlap for the top 5% of vertices exceeded chance levels, we conducted permutation testing for each parcel and participant, computing Bayes factors (BF10) (**Figure 7**). Under standard guidelines, BF10 values of 0-3 indicate anecdotal evidence, 3-10 moderate evidence, 10-30 strong evidence, 30-100 very strong evidence, and values greater than 100 extremely strong evidence in favor of the alternative hypothesis (i.e., greater overlap than expected by chance).

**Figure 7.**
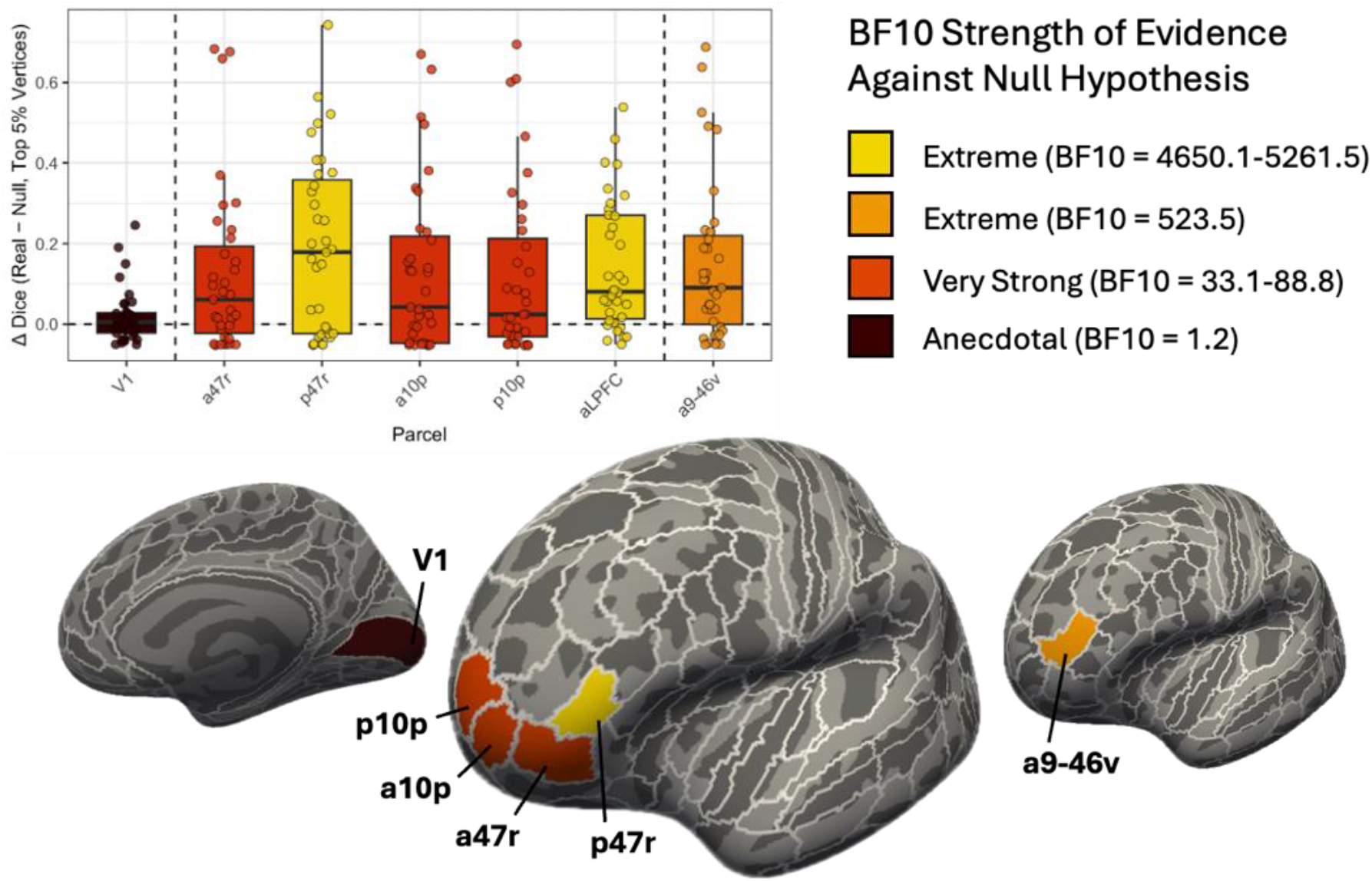
Degree of overlap between tasks varies across individuals and parcels in left aLPFC. Colors correspond to levels of BF10 magnitude as stated in Kelter (2020). To emphasize differences between extremely large BF10 values, the level of “Extreme” is split into two levels, with aLPFC as a whole showing the highest value numerically, followed by p47r and then benchmark region a9-46v. No parcels of interest showed Weak or Strong levels, and thus these levels are not included here. Brains feature V1 (left), individual aLPFC parcels (middle), and a9-46v (right). The broader aLPFC parcel comprises a47r, p47r, a10p, and p10p.

All four aLPFC parcels, as well as the broader aLPFC region, showed very strong to extremely strong evidence against the null hypothesis of random overlap (Kelter, 2020). Specifically, p47r (BF10 = 4650.06) showed extremely strong evidence, whereas a10p (BF10 = 88.83), p10p (BF10 = 33.09), and a47r (BF10 = 58) showed very strong evidence. By comparison, a9-46v showed extremely strong evidence against the null hypothesis (BF10 = 523.45), whereas V1 showed anecdotal evidence against it (BF10 = 1.20). The broader aLPFC region exhibited extremely strong evidence with a Bayes factor that was numerically far higher than the individual parcels (BF10 = 5261.46), which we anticipated because individuals’ clusters did not consistently fall into the same parcels. In sum, overlap between Reasoning and Memory tasks in aLPFC was significantly greater than expected by chance. However, the magnitude of overlap varied widely across individuals; this was true even when considering the whole aLPFC parcel.

##### 3.2.2.3 Exploratory correlations with behavior

Having established that clusters from the reasoning contrast were sensitive to the Memory manipulation, we tested whether activation on the Memory task in these clusters were meaningfully related to memory performance (d’) in the two parcels within which a substantial proportion of participants had significant clusters, a47r and p47r, as well as the broader aLPFC region. We found a significant positive correlation between activation and d’ for one of these three parcel definitions: p47r (**Figure 8A**). Although it did not survive false discovery rate (FDR) correction across the three analyses, the correlation remained stable under leave-one-subject-out jackknife resampling (full-sample *r* = 0.54), yielding correlations ranging from 0.42 to 0.62 across resamples.

**Figure 8:**
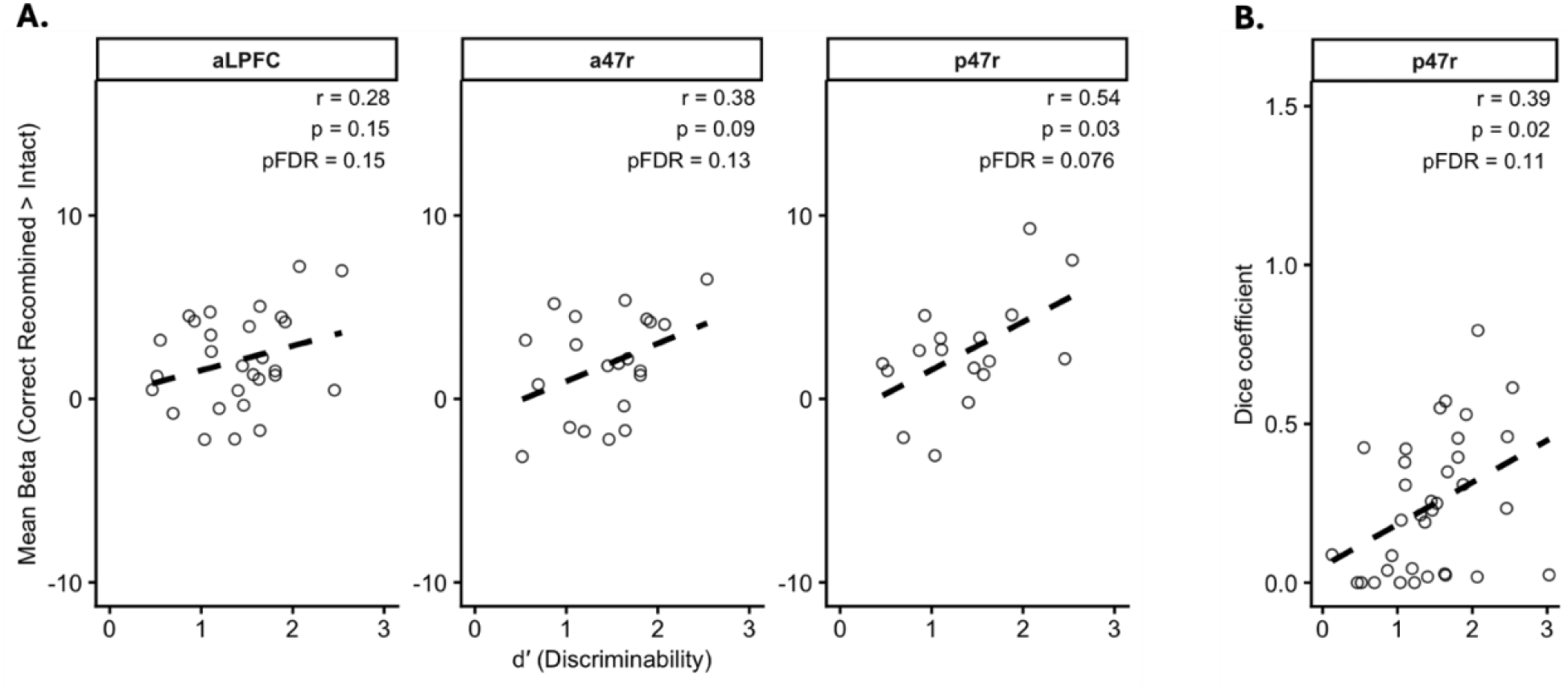
Memory performance associations with Memory task activation and vertex-level overlap between tasks. Each point represents one participant. Correlation coefficients (r), uncorrected p-values, and FDR-corrected p-values (pFDR) values are shown for each parcel. (A) Scatterplots show the relationship between recognition memory discriminability (d′) and the cluster-weighted mean beta within each parcel definition. (B) Scatterplot shows the relationship between recognition memory discriminability (d′) and Dice coefficients quantifying overlap between reasoning- and memory-related activation patterns (top 5% of vertices) within parcel p47r. Note that correlations between d’ and Dice coefficients were conducted for all four parcels of interest and the aLPFC parcel definition, and none were significant after FDR correction. Parcel p47r, shown here, was the only significant parcel at the uncorrected threshold.

In a second exploratory analysis, we tested whether the Dice coefficient for overlap between the top 5% of vertices engaged on the Reasoning and Memory tasks was associated with d’ on the Memory task (**Figure 8B**). Here, we considered all four individual aLPFC parcels (in addition to the broader aLPFC), because spatially contiguous activation across vertices was not essential for this analysis. Again here, as with the cluster analysis, only p47r showed a significant positive correlation. This correlation did not survive FDR correction across all five parcel definitions (or even when restricted to the four individual parcels: pFDR = .09); however, it was stable under leave-one-subject-out jackknife resampling (full-sample *r* = 0.39), yielding correlations ranging from 0.33 to 0.52 across resamples. In summary, both cluster activation and vertex overlap analyses tentatively implicated p47r in memory performance.

##### 3.2.2.4 Right hemisphere

As predicted, thereby motivating our planned focus on the left hemisphere, the prevalence of reasoning-related clusters was lower in the right hemisphere (see **Supplementary Figure 2**), and cluster-level sensitivity to memory was numerically reduced relative to the left-hemisphere regions (**Supplementary Figure 5A**). For comparison with the left hemisphere, we examined right hemisphere p47r and a47r, as well as the broader aLPFC even though only 23% of participants exhibited clusters in p47r (with 49% in a47r). Despite this reduced prevalence, Bayes factor analyses for participants exhibiting activation clusters provided evidence for greater overlap than expected by chance, albeit weaker than in the left hemisphere. Moreover, neither cluster-level betas nor Dice coefficients were correlated with d’ in the right hemisphere. Correlations between cluster-level betas and d’ ranged from r = 0.09 to 0.4 (all pFDR = 0.7). Similarly, associations between vertex-level overlap and d’ ranged from r = 0.05 to 0.32, with no effects surviving FDR correction (pFDR = 0.29-0.78) (**Supplementary Figure 7**). Taken together, right-hemisphere aLPFC regions exhibited weaker task overlap than their left-hemisphere counterparts, and showed no relationship to memory performance.

#### 3.2.3 Vis-What task analyses

To test whether the strength of overlap could be driven by the Reasoning task whose content is the most similar to the Memory task, as well as to more closely match the amount of data included for the two paradigms, we additionally conducted the same set of analyses replacing the average of all four Reasoning tasks with the Vis-What task. The conjunction and disjunction analyses revealed overlap patterns comparable to those observed across the full set of Reasoning tasks (**Supplementary Figure 9**). Cluster prevalence for the single Reasoning task was lower than for all four tasks, likely because of reduced statistical power, although the pattern across parcels was similar (**Supplementary Figure 10**). However, even with only one quarter of the Reasoning task data, 80% of participants had clusters somewhere in aLPFC – i.e., nearly as high as the prevalence rates for the full dataset. Sensitivity to the Memory manipulation was numerically weaker for Vis-What than across all Reasoning tasks, but was still significant for p47r and a47r (**Supplementary Figure 11**). Dice overlap at the vertex-level showed a somewhat different pattern across parcels when considering Vis-What in isolation (**Supplementary Figure 12**). For p47r, although the Bayes factor was still considered extremely strong for Vis-What than when all four tasks were included, it was numerically weaker (BF10 = 132 as compared with BF10 = 4650). On the other hand, evidence was stronger for Vis-What than for all four tasks in a47r and a10p, along with a9-46v. Thus, the degree of overlap observed in p47r for all tasks could not have been driven by the task with the most similar content (Vis-What); by contrast, shared demands for semantic processing of visually presented stimuli may have boosted overlap in other parcels.

Correlations with memory performance were observed in p47r for Vis-What alone, although the magnitudes differed numerically. For reasoning-related cluster-level mean betas, the correlation with d′ was stronger in p47r when analyses were restricted to the Vis-What task (r = .64; p-uncorrected = .01; **Supplementary Figure 13**) as compared to all four tasks (r = .54; p-uncorrected < .05), and was stable under leave-one-out jackknife resampling, yielding correlations ranging from 0.60 to 0.68. Further, the relationship for Vis-What survived when correcting for multiple comparisons across three parcel definitions (pFDR = .03), unlike the analysis involving all tasks (pFDR =.08). With regards to Dice coefficients, on the other hand, the correlation with d′ was similar in p47r for Vis-What (r = .38; p-uncorrected = .02) as for all four tasks (r = .39; p-uncorrected = .02), and was stable under leave-one-out jackknife resampling, yielding correlations ranging from 0.30 to 0.48 across resamples. However, as was the case across all Reasoning tasks, the correlation for p47r did not survive correction for multiple comparisons across all five parcel definitions (or even when correcting across only the four individual parcels: pFDR = .09 for Vis-What). Thus, while the cluster-level correlation was stronger for Vis-What, the relationship between d’ and overlap of the top 5% of vertices was not driven by any one of the four tasks.

## 4 DISCUSSION

In the present study, we sought to evaluate the possibility that aLPFC supports a foundational neurocognitive process that is shared across retrieval monitoring and relational thinking. The premise of this study is that evidence of neural overlap would be necessary, albeit not sufficient, for positing a shared cognitive function. However, aLPFC is a large, ill-defined area with heterogeneous task activations across participants – a fact that has often been overlooked in prior theorizing about aLPFC function (but see Reynolds et al., 2006; Westphal et al., 2016). As a preliminary step towards testing for a unifying role of aLPFC across multiple cognitive domains, we tested for overlap in fMRI activation across an associative memory retrieval monitoring task and four relational reasoning tasks involving images and sounds with varied stimulus content.

To assess degree of overlap at the group level, we performed a conjunction analysis that revealed overlapping activation within all aLPFC Glasser parcels, a47r, p47r, a10p, and p10p. Overlap was also observed in a number of more posterior LPFC regions, including but not limited to DLPFC regions 9-46d and 46 and VLPFC region IFSa, as well as DLPFC regions previously characterized as core MDS parcels: a9-46v and p9-46v. In contrast, disjunction analyses revealed only sparse, scattered vertices within aLPFC parcels, despite more robust reasoning-selective activation in anterior core MDS region a9-46v – along with memory-selective activation in left IFSa that likely reflects the fact that the Memory task involved words whereas three of the four Reasoning tasks were non-semantic. Thus, in keeping with Westphal et al. (2016), who showed widespread common activation across LPFC with no task differences in the left hemisphere, there were more commonalities than differences between tasks.

To test whether these reasoning-related regions were also sensitive to retrieval monitoring demands, we examined activation during the Memory task within clusters identified in individual participants. 80% of participants exhibited significant reasoning-related clusters at the predefined statistical threshold within left aLPFC, consistent with prior work showing engagement of this broader region in 10/10 participants on a relational reasoning localizer task (Smith et al., 2007). Notably, while these clusters were not cleanly confined to individual Glasser parcels, they were most frequently observed in parcels a47r and p47r. Clusters within these parcels, along with the broader aLPFC area, showed stronger activation for conditions with higher retrieval monitoring demands, suggesting that regions recruited during relational reasoning may also contribute to post-retrieval monitoring processes.

At a finer spatial scale, Bayesian analyses provided extremely strong evidence for overlap in spatial patterns of activation across tasks in p47r and aLPFC as a whole, along with very strong evidence in the other aLPFC parcels, even under a stringent criterion focusing on the top 5% of active vertices; this is notable given that the tasks differed from one another in multiple ways. Our benchmark regions contextualize the strength of these findings. There was only anecdotal evidence of overlap in a perceptually driven control region, V1. At the other end of the spectrum, a parcel widely implicated in cognitively demanding tasks, a9-46v, showed extremely strong evidence; notably, however, its BF10 value was not quite as high numerically as for p47r. Despite the striking results at the group level, a notable feature of the present results is the substantial variability in overlap across individuals. That is, although aggregate Bayesian evidence showed extremely or very strong evidence in favor of the alternative hypothesis (i.e., greater overlap than expected by chance) in aLPFC, some participants exhibited little or no overlap within a given parcel, or even across the broader aLPFC region.

Thus, we explored whether variability in neural overlap was behaviorally meaningful: specifically, whether it was related to memory performance, as reasoning performance was near ceiling. First, we found, for individually defined Reasoning task clusters in p47r, that stronger activation for the retrieval monitoring manipulation was positively associated with memory discrimination (d′). Second, we found that greater spatial correspondence (i.e., higher Dice coefficients) between the most 5% active vertices in p47r for each task was associated with d’. While these correlations did not survive correction for multiple comparisons, the fact that the same parcel was implicated in both types of analyses, while also being the one showing the strongest Dice overlap, all paints a coherent picture. That is, these results highlight p47r as a particularly promising candidate for shared computations supporting relational thinking and memory retrieval. We hypothesize that the degree of neural overlap in this area reflects the degree of engagement in relational thinking, or another shared neurocognitive process, during memory retrieval.

The implication of p47r across analyses is particularly noteworthy in relation to Westphal et al. (2016). In that study, a multi-voxel pattern analysis (MVPA) classifier trained to discriminate between analogical reasoning and source memory judgments revealed the peak region of task discriminability, at the group level, at MNI coordinates (−42, 46, 6). When mapped onto the Glasser multimodal parcellation, a spherical ROI of 6mm radius centered on this peak coordinate falls primarily within area p47r and extends into adjacent area a9-46v. When conducting an ROI analysis centered on this coordinate, Westphal and colleagues found that it was sensitive to both relational integration demands and source memory judgments. Thus, roughly the same region showing maximal neural overlap in the present study showed both maximal task discrimination as well as sensitivity to both task manipulations in this previous study. Because our Reasoning and Memory tasks were acquired in separate runs and differed substantially in task structure (blocked vs. event-related designs), we did not apply MVPA, as above-chance classification accuracy could reflect these low-level differences rather than distinct cognitive processes.

Broadly, the vertex-level Dice overlap in the present study can be reconciled with the prior MVPA analysis when one considers that our analyses focused on whether the same cortical vertices were preferentially recruited across tasks, whereas MVPA is sensitive to differences in distributed activation patterns, including the relative weighting of activity across shared neural populations. Thus, a region may recruit a common set of active vertices/voxels for two task manipulations while still exhibiting distinct patterns of strength and/or directionality of activation across tasks that allow a classifier to discriminate between them. We cannot explore this possibility further since, as noted above, our study design did not lend itself to MVPA. However, this interpretation is consistent with recent work demonstrating that a brain region clearly implicated in a particular cognitive function can nonetheless show distinguishable activity patterns between tasks that tap into this function – while not necessarily individuating tasks that do not (see results for inferior frontal gyrus and frontal eye fields in Chiou et al., 2024). Thus, overlap in recruitment and discriminability of activation patterns capture complementary aspects of neural organization.

Our findings also converge with several independent lines of evidence implicating left p47r, among other areas, in relational thinking. Peak coordinates from Neurosynth meta-analytic maps for the term “reasoning” or “relational”, as well as findings based on the multiple-demand system framework (Assem et al., 2020), both point to p47r as preferentially engaged during relational reasoning. Although one might expect this left inferior frontal lobe region in pars orbitalis to be preferentially involved in language processing, as in a recent study distinguishing between language, logic, and MDS systems (Kean et al., 2026), we found that clusters in p47r were recruited similarly on semantic and spatial tasks. At the same time, p47r lies posterior to the region that is most consistently referred to as anterior PFC or lateral frontal pole – although some publications (e.g., Baker et al., 2018) have included this parcel in the frontal pole, speaking to the importance of precise terminology. Moreover, although we are highlighting p47r in particular, it is worth mentioning that task-evoked activation did not neatly respect the Glasser parcel boundaries. This may be because these atlas regions are based on a fixed, group-average brain map that wasn’t tailored to our data, or because the true functional organization for these tasks is more nuanced than what current brain parcellation schemes capture.

Future work could investigate whether individual-specific anatomical or functional landmarks better predict activation patterns, or whether these regions exhibit a more flexible and task-dependent organization. More broadly, this line of work has the potential to refine theories of aLPFC function – or, at a minimum, of left aLPFC, given the possibility of hemispheric differences. Several important directions for future research remain. First, although the present study demonstrated robust neural overlap at the group level, substantial variability in the degree of overlap was observed across individuals, which may reflect meaningful differences in cognitive strategy or efficiency. Our exploratory findings suggest that greater neural overlap in p47r may be associated with better memory performance, but these brain-behavior relationships require replication in larger samples before firm conclusions can be drawn. Second, given initial evidence by Westphal et al. (2016) of differential functional connectivity between this area and large-scale resting-state networks as a function of task demands, future research could examine this more closely at the individual level. Third, the present findings provide a principled target for future studies to evaluate whether this same region is recruited across other functions that have been attributed to aLPFC, including planning, prospective memory, exploration, and decision-making. Such work could help distinguish between competing accounts of aLPFC function by determining which cognitive operations consistently recruit this region across diverse task contexts, and via which patterns of coordinated activity across the brain.

## Data and Code Availability

Available upon request.

## Author Contributions

**S.A.B.:** Manuscript writing, Manuscript review & editing, Supervision, Project administration, Investigation, Funding acquisition, Conceptualization. **A.C.:** Manuscript writing, Manuscript review & editing, Project administration, Visualizations, Methodology, Investigation, Formal analysis, Data curation, Conceptualization. **M.V.:** Manuscript review & editing, Methodology, Investigation, Data curation, Conceptualization. **M.A.:** Manuscript review & editing, Supervision, Methodology.

## Declaration of the use of AI

AI was used to check spelling and grammar, as well as recommend potential analyses.

## Acknowledgements

We thank Jasper Liu, Jacquelyn Borcea, Narod Berberian, Sarah Peykar, and Suvi Häkkinen for their efforts in initial piloting and preparation of this project. We also thank Natalie Kwak, Sarah Peykar, and Shirley Nguyen for their assistance with data collection. Lastly, we thank Nick Ichien for providing code for stimulus construction of the memory task.

## REFERENCES

Abraham, A., Pedregosa, F., Eickenberg, M., Gervais, P., Mueller, A., Kossaifi, J., Gramfort, A., Thirion, B., & Varoquaux, G. (2014). Machine learning for neuroimaging with scikit-learn. Frontiers in Neuroinformatics, 8, Article 14.

Achim, A. M., & Lepage, M. (2005). Dorsolateral prefrontal cortex involvement in memory post-retrieval monitoring revealed in both item and associative recognition tests. NeuroImage, 24(4), 1113–1121. 10.1016/j.neuroimage.2004.10.036

Alexander, P. A. (2016). Relational thinking and relational reasoning: Harnessing the power of patterning. npj Science of Learning, 1, Article 16004. 10.1038/npjscilearn.2016.4

Altmayer, V., Ovando-Tellez, M., Bieth, T., Batrancourt, B., Rametti-Lacroux, A., Moreno-Rodriguez, S., Bouzigues, A., Ledu, V., Garcin, B., Lopez-Persem, A., Margulies, D., Levy, R., Volle, E., & ECOCAPTURE study group. (2026). A rostral prefrontal mediolateral gradient predicts creativity in frontotemporal dementia. Brain, awag032. 10.1093/brain/awag032

Andersson, J. L. R., Skare, S., & Ashburner, J. (2003). How to correct susceptibility distortions in spin-echo echo-planar images: Application to diffusion tensor imaging. NeuroImage, 20(2), 870– 888.

Assem, M., Glasser, M. F., Van Essen, D. C., & Duncan, J. (2020). A Domain-General Cognitive Core Defined in Multimodally Parcellated Human Cortex. Cerebral cortex (New York, N.Y. : 1991), 30(8), 4361–4380. 10.1093/cercor/bhaa023

Assem, M., Shashidhara, S., Glasser, M. F., & Duncan, J. (2022). Precise Topology of Adjacent Domain-General and Sensory-Biased Regions in the Human Brain. Cerebral cortex (New York, N.Y. : 1991), 32(12), 2521–2537. 10.1093/cercor/bhab362

Avants, B. B., Epstein, C. L., Grossman, M., & Gee, J. C. (2008). Symmetric diffeomorphic image registration with cross-correlation: Evaluating automated labeling of elderly and neurodegenerative brain. Medical Image Analysis, 12(1), 26–41.

Azuar, C., Reyes, P., Slachevsky, A., Volle, E., Kinkingnehun, S., Kouneiher, F., Bravo, E., Dubois, B., Koechlin, E., & Levy, R. (2014). Testing the model of caudo-rostral organization of cognitive control in the human with frontal lesions. NeuroImage, 84, 1053–1060. 10.1016/j.neuroimage.2013.09.031

Badre, D., & D’Esposito, M. (2007). Functional magnetic resonance imaging evidence for a hierarchical organization of the prefrontal cortex. Journal of cognitive neuroscience, 19(12), 2082– 2099. 10.1162/jocn.2007.19.12.2082

Badre, D., & D’Esposito, M. (2009). Is the rostro-caudal axis of the frontal lobe hierarchical? Nature Reviews Neuroscience, 10(9), 659–669. 10.1038/nrn2667

Bahlmann, J., Aarts, E., & D’Esposito, M. (2015). Influence of motivation on control hierarchy in the human frontal cortex. The Journal of neuroscience : the official journal of the Society for Neuroscience, 35(7), 3207–3217. 10.1523/JNEUROSCI.2389-14.2015

Baird, B., Smallwood, J., Gorgolewski, K. J., & Margulies, D. S. (2013). Medial and lateral networks in anterior prefrontal cortex support metacognitive ability for memory and perception. The Journal of neuroscience : the official journal of the Society for Neuroscience, 33(42), 16657– 16665. 10.1523/JNEUROSCI.0786-13.2013

Baker, C. M., Burks, J. D., Briggs, R. G., Conner, A. K., Glenn, C. A., Morgan, J. P., Stafford, J., Sali, G., McCoy, T. M., Battiste, J. D., O’Donoghue, D. L., & Sughrue, M. E. (2018). A Connectomic Atlas of the Human Cerebrum-Chapter 2: The Lateral Frontal Lobe. Operative neurosurgery (Hagerstown, Md.), 15(suppl_1), S10–S74. 10.1093/ons/opy254

Behzadi, Y., Restom, K., Liau, J., & Liu, T. T. (2007). A component based noise correction method (CompCor) for BOLD and perfusion based fMRI. NeuroImage, 37(1), 90–101.

Bendetowicz, D., Urbanski, M., Garcin, B., Foulon, C., Levy, R., Bréchemier, M. L., Rosso, C., Thiebaut de Schotten, M., & Volle, E. (2018). Two critical brain networks for generation and combination of remote associations. Brain : a journal of neurology, 141(1), 217–233. 10.1093/brain/awx294

Bludau, S., Eickhoff, S. B., Mohlberg, H., Caspers, S., Laird, A. R., Fox, P. T., Schleicher, A., Zilles, K., & Amunts, K. (2014). Cytoarchitecture, probability maps and functions of the human frontal pole. NeuroImage, 93 Pt 2(Pt 2), 260–275. 10.1016/j.neuroimage.2013.05.052

Blumenfeld, R. S., & Ranganath, C. (2019). The lateral prefrontal cortex and human long-term memory. Handbook of clinical neurology, 163, 221–235. 10.1016/B978-0-12-804281-6.00012-4

Boorman, E. D., Behrens, T. E., Woolrich, M. W., & Rushworth, M. F. (2009). How green is the grass on the other side? Frontopolar cortex and the evidence in favor of alternative courses of action. Neuron, 62(5), 733–743. 10.1016/j.neuron.2009.05.014

Boschin, E., Ainsworth, M., Galeazzi, J., & Buckley, M. (2025). Memories or Decisions? Bridging Accounts of Frontopolar Function. Neuropsychologia. 211. 109119. 10.1016/j.neuropsychologia.2025.109119.

Braver, T. S., & Bongiolatti, S. R. (2002). The role of frontopolar cortex in subgoal processing during working memory. NeuroImage, 15(3), 523–536. 10.1006/nimg.2001.1019

Brysbaert, M., Warriner, A.B. & Kuperman, V. (2014). Concreteness ratings for 40 thousand generally known English word lemmas. Behav Res 46, 904–911. 10.3758/s13428-013-0403-5

Buckner, R. L., Koutstaal, W., Schacter, D. L., Dale, A. M., Rotte, M., & Rosen, B. R. (1998). Functional-anatomic study of episodic retrieval. II. Selective averaging of event-related fMRI trials to test the retrieval success hypothesis. NeuroImage, 7(3), 163–175. 10.1006/nimg.1998.0328

Bunge, S. A., Wendelken, C., Badre, D., & Wagner, A. D. (2005). Analogical reasoning and prefrontal cortex: evidence for separable retrieval and integration mechanisms. Cerebral cortex (New York, N.Y. : 1991), 15(3), 239–249. 10.1093/cercor/bhh126

Bunge, S. A., & Zelazo, P. D. (2006). A brain-based account of the development of rule use in childhood. Current Directions in Psychological Science, 15(3), 118–121. 10.1111/j.0963-7214.2006.00419.x

Bunge, S. A., & Wendelken, C. (2009). Comparing the bird in the hand with the ones in the bush. Neuron, 62(5), 609–611. 10.1016/j.neuron.2009.05.020

Bunge, S. A., Helskog, E. H., & Wendelken, C. (2009). Left, but not right, rostrolateral prefrontal cortex meets a stringent test of the relational integration hypothesis. NeuroImage, 46(1), 338– 342. 10.1016/j.neuroimage.2009.01.064

Burgess, P. W., Veitch, E., de Lacy Costello, A., & Shallice, T. (2000). The cognitive and neuroanatomical correlates of multitasking. Neuropsychologia, 38(6), 848–863. 10.1016/s0028-3932(99)00134-7

Burgess, P. Strategy application disorder: the role of the frontal lobes in human multitasking. Psychological Research Psychologische Forschung 63, 279–288 (2000). 10.1007/s004269900006

Burgess, P. W., Scott, S. K., & Frith, C. D. (2003). The role of the rostral frontal cortex (area 10) in prospective memory: A lateral versus medial dissociation. Neuropsychologia, 41(8), 906–918. 10.1016/S0028-3932(02)00327-5

Burgess, P. W., Dumontheil, I., & Gilbert, S. J. (2007). The gateway hypothesis of rostral prefrontal cortex (area 10) function. Trends in cognitive sciences, 11(7), 290–298. 10.1016/j.tics.2007.05.004

Cabeza, R., & Nyberg, L. (2000). Imaging cognition II: An empirical review of 275 PET and fMRI studies. Journal of cognitive neuroscience, 12(1), 1–47. 10.1162/08989290051137585

Christoff, K., Gabrieli, J.D.E. (2000). The frontopolar cortex and human cognition: Evidence for a rostrocaudal hierarchical organization within the human prefrontal cortex. Psychobiology 28, 168– 186. 10.3758/BF03331976

Christoff, K., Prabhakaran, V., Dorfman, J., Zhao, Z., Kroger, J. K., Holyoak, K. J., & Gabrieli, J. D. (2001). Rostrolateral prefrontal cortex involvement in relational integration during reasoning. NeuroImage, 14(5), 1136–1149. 10.1006/nimg.2001.0922

Christoff, K., Keramatian, K., Gordon, A. M., Smith, R., & Mädler, B. (2009). Prefrontal organization of cognitive control according to levels of abstraction. Brain research, 1286, 94–105. 10.1016/j.brainres.2009.05.096

Ciric, R., Thompson, W. H., Lorenz, R., Goncalves, M., MacNicol, E., Markiewicz, C. J., Bhagwat, N., Sivaraju, V., Snider, K., Amlien, I. K., Feingold, F., Poldrack, R. A., Gorgolewski, K. J., & Esteban, O. (2022). TemplateFlow: FAIR-sharing of multi-scale, multi-species brain models. Nature Methods, 19(12), 1568–1571.

Dale, A. M., Fischl, B., & Sereno, M. I. (1999). Cortical surface-based analysis: I. Segmentation and surface reconstruction. NeuroImage, 9(2), 179–194.

Desrochers, T. M., Chatham, C. H., & Badre, D. (2015). The Necessity of Rostrolateral Prefrontal Cortex for Higher-Level Sequential Behavior. Neuron, 87(6), 1357–1368. 10.1016/j.neuron.2015.08.026

Dobbins, I. G., Foley, H., Schacter, D. L., & Wagner, A. D. (2002). Executive control during episodic retrieval: multiple prefrontal processes subserve source memory. Neuron, 35(5), 989– 996. 10.1016/s0896-6273(02)00858-9

Dobbins, I. G., & Han, S. (2006). Isolating rule-versus evidence-based prefrontal activity during episodic and lexical discrimination: A functional magnetic resonance imaging investigation of detection theory distinctions. Cerebral Cortex, 16(11), 1614–1622. 10.1093/cercor/bhj098

Donahue CJ, Glasser MF, Preuss TM, Rilling JK, Van Essen DC. Quantitative assessment of prefrontal cortex in humans relative to nonhuman primates. Proc Natl Acad Sci U S A. 2018;115(22):E5183–E5192. doi:10.1073/pnas.1721653115

Duarte, A., Ranganath, C., & Knight, R. T. (2005). Effects of unilateral prefrontal lesions on familiarity, recollection, and source memory. Journal of Neuroscience, 25(36), 8333–8337. 10.1523/JNEUROSCI.1392-05.2005

Duncan, J., Assem, M., & Shashidhara, S. (2020). Integrated Intelligence from Distributed Brain Activity. Trends in cognitive sciences, 24(10), 838–852. 10.1016/j.tics.2020.06.012

Dreher, J.-C., Koechlin, E., Tierney, M. & Grafman, J. Damage to the Fronto-Polar Cortex Is Associated with Impaired Multitasking. PLoS One 3, e3227 (2008)

Esteban, O., Blair, R., Markiewicz, C. J., et al. (2018). fMRIPrep [Software]. Zenodo. 10.5281/zenodo.852659

Esteban, O., Markiewicz, C. J., Blair, R. W., Moodie, C. A., Isik, A. I., Erramuzpe, A., Kent, J. D., Goncalves, M., DuPre, E., Snyder, M., Oya, H., Ghosh, S. S., Wright, J., Durnez, J., Poldrack, R. A., & Gorgolewski, K. J. (2019). fMRIPrep: a robust preprocessing pipeline for functional MRI. Nature methods, 16(1), 111–116. 10.1038/s41592-018-0235-4

Fandakova, Y., Bunge, S. A., Wendelken, C., Desautels, P., Hunter, L., Lee, J. K., & Ghetti, S. (2018). The Importance of Knowing When You Don’t Remember: Neural Signaling of Retrieval Failure Predicts Memory Improvement Over Time. Cerebral cortex (New York, N.Y. : 1991), 28(1), 90–102. 10.1093/cercor/bhw352

Fedorenko, E., Duncan, J., & Kanwisher, N. (2013). Broad domain generality in focal regions of frontal and parietal cortex. Proceedings of the National Academy of Sciences of the United States of America, 110(41), 16616–16621. 10.1073/pnas.1315235110

Fleming, S. M., & Dolan, R. J. (2012). The neural basis of metacognitive ability. Philosophical transactions of the Royal Society of London. Series B, Biological sciences, 367(1594), 1338– 1349. 10.1098/rstb.2011.0417

Fleming, S. M., Huijgen, J., & Dolan, R. J. (2012). Prefrontal contributions to metacognition in perceptual decision making. The Journal of neuroscience : the official journal of the Society for Neuroscience, 32(18), 6117–6125. 10.1523/JNEUROSCI.6489-11.2012

Fleming, S. M., Ryu, J., Golfinos, J. G., & Blackmon, K. E. (2014). Domain-specific impairment in metacognitive accuracy following anterior prefrontal lesions. Brain : a journal of neurology, 137(Pt 10), 2811–2822. 10.1093/brain/awu221

Fonov, V., Evans, A. C., Botteron, K., Almli, C. R., McKinstry, R. C., Collins, D. L., & Brain Development Cooperative Group. (2011). Unbiased average age-appropriate atlases for pediatric studies. NeuroImage, 54(1), 313–327.

Fuster J. M. (2001). The prefrontal cortex--an update: time is of the essence. Neuron, 30(2), 319–333. 10.1016/s0896-6273(01)00285-9

Fuster J. M. (2004). Upper processing stages of the perception-action cycle. Trends in cognitive sciences, 8(4), 143–145. 10.1016/j.tics.2004.02.004

Gilbert, S. J., Spengler, S., Simons, J. S., Steele, J. D., Lawrie, S. M., Frith, C. D., & Burgess, P. W. (2006). Functional specialization within rostral prefrontal cortex (area 10): a meta-analysis. Journal of cognitive neuroscience, 18(6), 932–948. 10.1162/jocn.2006.18.6.932

Gilbert, S. J., Gonen-Yaacovi, G., Benoit, R. G., Volle, E., & Burgess, P. W. (2010). Distinct functional connectivity associated with lateral versus medial rostral prefrontal cortex: a meta-analysis. NeuroImage, 53(4), 1359–1367. 10.1016/j.neuroimage.2010.07.032

Glasser, M., Coalson, T., Robinson, E. et al. A multi-modal parcellation of human cerebral cortex. Nature 536, 171–178 (2016). 10.1038/nature18933

Gorgolewski, K., Burns, C. D., Madison, C., Clark, D., Halchenko, Y. O., Waskom, M. L., & Ghosh, S. S. (2011). Nipype: A flexible, lightweight and extensible neuroimaging data processing framework in Python. Frontiers in Neuroinformatics, 5, Article 13.

Gorgolewski, K. J., Auer, T., Calhoun, V. D., Craddock, R. C., Das, S., Duff, E. P., Flandin, G., Ghosh, S. S., Glatard, T., Halchenko, Y. O., Handwerker, D. A., Hanke, M., Keator, D., Li, X., Michael, Z., Maumet, C., Nichols, B. N., Nichols, T. E., Pellman, J., Poline, J.-B., Rokem, A., Schaefer, G., Sochat, V., Triplett, W., Turner, J. A., Varoquaux, G., & Poldrack, R. A. (2016). The brain imaging data structure, a format for organizing and describing outputs of neuroimaging experiments. Scientific Data, 3, 160044. 10.1038/sdata.2016.44

Gorgolewski, K. J., Esteban, O., Markiewicz, C. J., et al. (2018). Nipype [Software]. Zenodo.

Greve, D. N., & Fischl, B. (2009). Accurate and robust brain image alignment using boundary-based registration. NeuroImage, 48(1), 63–72.

Grinband, J., Wager, T. D., Lindquist, M., Ferrera, V. P., & Hirsch, J. (2008). Detection of time-varying signals in event-related fMRI designs. NeuroImage, 43(3), 509–520. 10.1016/j.neuroimage.2008.07.065

Halford, G. S., Wilson, W. H., & Phillips, S. (2010). Relational knowledge: the foundation of higher cognition. Trends in cognitive sciences, 14(11), 497–505. 10.1016/j.tics.2010.08.005

Henson, R. N., Shallice, T., & Dolan, R. J. (1999). Right prefrontal cortex and episodic memory retrieval: a functional MRI test of the monitoring hypothesis. Brain : a journal of neurology, 122 ( *Pt 7*), 1367–1381. 10.1093/brain/122.7.1367

Henson, R. N., Rugg, M. D., Shallice, T., Josephs, O., & Dolan, R. J. (1999). Recollection and familiarity in recognition memory: an event-related functional magnetic resonance imaging study. The Journal of neuroscience : the official journal of the Society for Neuroscience, 19(10), 3962– 3972. 10.1523/JNEUROSCI.19-10-03962.1999

Hobeika, L., Diard-Detoeuf, C., Garcin, B., Levy, R., & Volle, E. (2016). General and specialized brain correlates for analogical reasoning: A meta-analysis of functional imaging studies. Human brain mapping, 37(5), 1953–1969. 10.1002/hbm.23149

Hogeveen, J., Medalla, M., Ainsworth, M., Galeazzi, J. M., Hanlon, C. A., Mansouri, F. A., & Costa, V. D. (2022). What does the frontopolar cortex contribute to goal-directed cognition and action? Journal of Neuroscience, 42(45), 8508–8513. 10.1523/JNEUROSCI.1herewith2666-22.2022

Holyoak, K. J., & Monti, M. M. (2021). Relational Integration in the Human Brain: A Review and Synthesis. Journal of cognitive neuroscience, 33(3), 341–356. 10.1162/jocn_a_01619

Ichien, N., Lu, H., & Holyoak, K. J. (2022). Predicting patterns of similarity among abstract semantic relations. Journal of Experimental Psychology: Learning, Memory, and Cognition, 48(1), 108–121. 10.1037/xlm0001010

Ichien, N., Alfred, K. L., Baia, S., Kraemer, D. J. M., Holyoak, K. J., Bunge, S. A., & Lu, H. (2023). Relational and lexical similarity in analogical reasoning and recognition memory: Behavioral evidence and computational evaluation. Cognitive psychology, 141, 101550. 10.1016/j.cogpsych.2023.101550

Jenkinson, M., Bannister, P., Brady, M., & Smith, S. (2002). Improved optimization for the robust and accurate linear registration and motion correction of brain images. NeuroImage, 17(2), 825– 841.

Kean, H., Fung, A., Jaggers, P., Chen, J., Rule, J. S., Benn, Y., Tenenbaum, J. B., Piantadosi, S. T., Varley, R. A., & Fedorenko, E. (2026). Evidence from formal logical reasoning reveals that the language of thought is not natural language. Proceedings of the National Academy of Sciences, 123(28), e2520095123. 10.1073/pnas.2520095123

Kelly, P. G., Bortfeld, H., Bunge, S. A., & Joyner, K. J. (2026). Evaluating the reliability of functional near-infrared spectroscopy data in the context of a reasoning paradigm. Developmental Cognitive Neuroscience, 80, 101763. 10.1016/j.dcn.2026.101763

Kelter R. (2020). Bayesian alternatives to null hypothesis significance testing in biomedical research: a non-technical introduction to Bayesian inference with JASP. BMC medical research methodology, 20(1), 142. 10.1186/s12874-020-00980-6

Klein, A., Ghosh, S. S., Bao, F. S., Giard, J., Häme, Y., Stavsky, E., Lee, N., Rossa, B., Reuter, M., Neto, E. C., & Keshavan, A. (2017). Mindboggling morphometry of human brains. PLOS Computational Biology, 13(2), Article e1005350.

Knowlton, B. J., Morrison, R. G., Hummel, J. E., & Holyoak, K. J. (2012). A neurocomputational system for relational reasoning. Trends in Cognitive Sciences, 16(7), 373–381. 10.1016/j.tics.2012.06.002

Koechlin, E., Basso, G., Pietrini, P., Panzer, S., & Grafman, J. (1999). The role of the anterior prefrontal cortex in human cognition. Nature, 399(6732), 148–151. 10.1038/20178

Koechlin, E., Ody, C., & Kouneiher, F. (2003). The architecture of cognitive control in the human prefrontal cortex. Science (New York, N.Y.), 302(5648), 1181–1185. 10.1126/science.1088545

Koechlin, E., & Hyafil, A. (2007). Anterior prefrontal function and the limits of human decision-making. Science (New York, N.Y.), 318(5850), 594–598. 10.1126/science.1142995

Krawczyk D. C. (2012). The cognition and neuroscience of relational reasoning. Brain research, 1428, 13–23. 10.1016/j.brainres.2010.11.080

Kroger, J. K., Sabb, F. W., Fales, C. L., Bookheimer, S. Y., Cohen, M. S., & Holyoak, K. J. (2002). Recruitment of anterior dorsolateral prefrontal cortex in human reasoning: a parametric study of relational complexity. Cerebral cortex, 12(5), 477–485.

Lepage, M., Ghaffar, O., Nyberg, L., & Tulving, E. (2000). Prefrontal cortex and episodic memory retrieval mode. Proceedings of the National Academy of Sciences of the United States of America, 97(1), 506–511. 10.1073/pnas.97.1.506

Liu H, Qin W, Li W, Fan L, Wang J, Jiang T, Yu C (2013) Connectivity-based parcellation of the human frontal pole with diffusion tensor imaging. JNeurosci 33:6782–6790

Lu, H., Chen, D., & Holyoak, K. J. (2012). Bayesian analogy with relational transformations. Psychological review, 119(3), 617–648. 10.1037/a0028719

Mansouri, F. A., Koechlin, E., Rosa, M. G. P., & Buckley, M. J. (2017). Managing competing goals -a key role for the frontopolar cortex. Nature reviews. Neuroscience, 18(11), 645–657. 10.1038/nrn.2017.111

Mars, R. B., Jbabdi, S., Sallet, J., O’Reilly, J. X., Croxson, P. L., Olivier, E., Noonan, M. P., Bergmann, C., Mitchell, A. S., Baxter, M. G., Behrens, T. E., Johansen-Berg, H., Tomassini, V., Miller, K. L., & Rushworth, M. F. (2011). Diffusion-weighted imaging tractography-based parcellation of the human parietal cortex and comparison with human and macaque resting-state functional connectivity. The Journal of neuroscience : the official journal of the Society for Neuroscience, 31(11), 4087–4100. 10.1523/JNEUROSCI.5102-10.2011

Moayedi, M., Salomons, T. V., Dunlop, K. A., Downar, J., & Davis, K. D. (2015). Connectivity-based parcellation of the human frontal polar cortex. Brain structure & function, 220(5), 2603– 2616. 10.1007/s00429-014-0809-6

Morales, J., Lau, H., & Fleming, S. M. (2018). Domain-general and domain-specific patterns of activity supporting metacognition in human prefrontal cortex. Journal of Neuroscience, 38(14), 3534–3546. 10.1523/JNEUROSCI.2360-17.2018

Nyberg, L., Tulving, E., Habib, R., Nilsson, L.-G., Kapur, S., Houle, S., Cabeza, R., & McIntosh, A. R. (1995). Functional brain maps of retrieval mode and recovery of episodic information. NeuroReport, 7(1), 249–252.

Orr, J. M., Smolker, H. R., & Banich, M. T. (2015). Organization of the human frontal pole revealed by large-scale DTI-based connectivity: Implications for control of behavior. PLOS ONE, 10(5), e0124797. 10.1371/journal.pone.0124797

Patriat, R., Reynolds, R. C., & Birn, R. M. (2017). An improved model of motion-related signal changes in fMRI. NeuroImage, 144(Pt A), 74–82.

Popov, V., Hristova, P., & Anders, R. (2017). The relational luring effect: Retrieval of relational information during associative recognition. Journal of experimental psychology. General, 146(5), 722–745. 10.1037/xge0000305

Power, J. D., Mitra, A., Laumann, T. O., Snyder, A. Z., Schlaggar, B. L., & Petersen, S. E. (2014). Methods to detect, characterize, and remove motion artifact in resting state fMRI. NeuroImage, 84, 320–341.

Ramnani, N., & Owen, A. M. (2004). Anterior prefrontal cortex: insights into function from anatomy and neuroimaging. Nature reviews. Neuroscience, 5(3), 184–194. 10.1038/nrn1343

Ranganath, C., Johnson, M. K., & D’Esposito, M. (2000). Left anterior prefrontal activation increases with demands to recall specific perceptual information. The Journal of neuroscience : the official journal of the Society for Neuroscience, 20(22), RC108. 10.1523/JNEUROSCI.20-22-j0005.2000

Ranganath, C., & Paller, K. A. (2000). Neural correlates of memory retrieval and evaluation. Brain research. Cognitive brain research, 9(2), 209–222. 10.1016/s0926-6410(99)00048-8

Ranganath, C., Heller, A. S., & Wilding, E. L. (2007). Dissociable correlates of two classes of retrieval processing in prefrontal cortex. NeuroImage, 35(4), 1663–1673. 10.1016/j.neuroimage.2007.01.020

Reynolds, J. R., McDermott, K. B., & Braver, T. S. (2006). A direct comparison of anterior prefrontal cortex involvement in episodic retrieval and integration. *Cerebral cortex (New York*, N.Y*. :* 1991), *16*(4), 519–528. 10.1093/cercor/bhi131

Roca, M., Parr, A., Thompson, R., Woolgar, A., Torralva, T., Antoun, N., Manes, F., & Duncan, J. (2010). Executive function and fluid intelligence after frontal lobe lesions. Brain : a journal of neurology, 133(Pt 1), 234–247. 10.1093/brain/awp269

Roca, M., Torralva, T., Gleichgerrcht, E., Woolgar, A., Thompson, R., Duncan, J., & Manes, F. (2011). The role of Area 10 (BA10) in human multitasking and in social cognition: a lesion study. Neuropsychologia, 49(13), 3525–3531. 10.1016/j.neuropsychologia.2011.09.003

Rosa, M. G. P., Soares, J. G. M., Chaplin, T. A., Majka, P., Bakola, S., Phillips, K. A., Reser, D. H., & Gattass, R. (2019). Cortical Afferents of Area 10 in Cebus Monkeys: Implications for the Evolution of the Frontal Pole. *Cerebral cortex (New York*, N.Y*. :* 1991), *29*(4), 1473–1495. 10.1093/cercor/bhy044

Rugg, M. D., & Henson, R. N. A. (2002). Episodic memory retrieval: an (event-related) functional neuroimaging perspective. The Cognitive Neuroscience of True and False Memories, 3–37

Satterthwaite, T. D., Elliott, M. A., Gerraty, R. T., Ruparel, K., Loughead, J., Calkins, M. E., Eickhoff, S. B., Hakonarson, H., Gur, R. C., Gur, R. E., & Wolf, D. H. (2013). An improved framework for confound regression and filtering for control of motion artifact in the preprocessing of resting-state functional connectivity data. NeuroImage, 64, 240–256.

Semendeferi, K., Teffer, K., Buxhoeveden, D. P., Park, M. S., Bludau, S., Amunts, K., Travis, K., & Buckwalter, J. (2011). Spatial organization of neurons in the frontal pole sets humans apart from great apes. Cerebral Cortex, 21(7), 1485–1497. 10.1093/cercor/bhq191

Shekhar, M., & Rahnev, D. (2018). Distinguishing the Roles of Dorsolateral and Anterior PFC in Visual Metacognition. The Journal of neuroscience : the official journal of the Society for Neuroscience, 38(22), 5078–5087. 10.1523/JNEUROSCI.3484-17.2018

Simons, J. S., Henson, R. N., Gilbert, S. J., & Fletcher, P. C. (2008). Separable forms of reality monitoring supported by anterior prefrontal cortex. Journal of cognitive neuroscience, 20(3), 447– 457. 10.1162/jocn.2008.20036

Smith, R., Keramatian, K., & Christoff, K. (2007). Localizing the rostrolateral prefrontal cortex at the individual level. NeuroImage, 36(4), 1387–1396. 10.1016/j.neuroimage.2007.04.032

Spencer, D., Yue, Y. R., Bolin, D., Ryan, S., & Mejia, A. F. (2022). Simultaneous confidence regions for spatial excursion sets, with an application to fMRI data. Journal of Computational and Graphical Statistics, 31(3), 651–662. 10.1080/10618600.2021.2001379

Spunt, B. (2016). easy-optimize-x: Formal Release for Archiving on Zenodo (Version 1.0) [Computer software]. Zenodo. 10.5281/zenodo.58616

Starr, A., Leib, E. R., Younger, J. W., Project iLead Consortium, Uncapher, M. R., & Bunge, S. A. (2023). Relational thinking: An overlooked component of executive functioning. Developmental science, 26(3), e13320. 10.1111/desc.13320

Tranel, D., Manzel, K., & Anderson, S. W. (2008). Is the prefrontal cortex important for fluid intelligence? A neuropsychological study using Matrix Reasoning. The Clinical neuropsychologist, 22(2), 242–261. 10.1080/13854040701218410

Turner, M. S., Cipolotti, L., Yousry, T. A., & Shallice, T. (2008). Confabulation: damage to a specific inferior medial prefrontal system. Cortex; a journal devoted to the study of the nervous system and behavior, 44(6), 637–648. 10.1016/j.cortex.2007.01.002

Tustison, N. J., Avants, B. B., Cook, P. A., Zheng, Y., Egan, A., Yushkevich, P. A., & Gee, J. C. (2010). N4ITK: Improved N3 bias correction. IEEE Transactions on Medical Imaging, 29(6), 1310– 1320.

Underwood AG, Guynn MJ and Cohen A-L (2015). The Future Orientation of Past Memory: The Role of BA 10 in Prospective and Retrospective Retrieval Modes. Front. Hum. Neurosci. 9:668. doi: 10.3389/fnhum.2015.00668

Urbanski, M., Bréchemier, M. L., Garcin, B., Bendetowicz, D., Thiebaut de Schotten, M., Foulon, C., Rosso, C., Clarençon, F., Dupont, S., Pradat-Diehl, P., Labeyrie, M. A., Levy, R., & Volle, E. (2016). Reasoning by analogy requires the left frontal pole: lesion-deficit mapping and clinical implications. Brain : a journal of neurology, 139(Pt 6), 1783–1799. 10.1093/brain/aww072

Vashel M., Chen, A., Bunge, S. A. (in prep).

Vendetti, M. S., & Bunge, S. A. (2014). Evolutionary and developmental changes in the lateral frontoparietal network: a little goes a long way for higher-level cognition. Neuron, 84(5), 906–917. 10.1016/j.neuron.2014.09.035

Velanova, K., Jacoby, L. L., Wheeler, M. E., McAvoy, M. P., Petersen, S. E., & Buckner, R. L. (2003). Functional-anatomic correlates of sustained and transient processing components engaged during controlled retrieval. The Journal of neuroscience : the official journal of the Society for Neuroscience, 23(24), 8460–8470. 10.1523/JNEUROSCI.23-24-08460.2003

Wagner AD, Shannon BJ, Kahn I, Buckner RL (2005): Parietal lobe contributions to episodic memory retrieval. Trends Cogn Sci 9:445–453.

Wendelken, C., Nakhabenko, D., Donohue, S. E., Carter, C. S., & Bunge, S. A. (2008). “Brain is to thought as stomach is to ??”: investigating the role of rostrolateral prefrontal cortex in relational reasoning. Journal of cognitive neuroscience, 20(4), 682–693. 10.1162/jocn.2008.20055

Wendelken, C., Bunge, S. A., & Carter, C. S. (2008). Maintaining structured information: an investigation into functions of parietal and lateral prefrontal cortices. Neuropsychologia, 46(2), 665–678. 10.1016/j.neuropsychologia.2007.09.015

Wendelken, C., & Bunge, S. A. (2010). Transitive inference: distinct contributions of rostrolateral prefrontal cortex and the hippocampus. Journal of cognitive neuroscience, 22(5), 837–847. 10.1162/jocn.2009.21226

Wendelken, C., Chung, D., & Bunge, S. A. (2012). Rostrolateral prefrontal cortex: domain-general or domain-sensitive?. Human brain mapping, 33(8), 1952–1963. 10.1002/hbm.21336

Westphal, A. J., Reggente, N., Ito, K. L., & Rissman, J. (2016). Shared and distinct contributions of rostrolateral prefrontal cortex to analogical reasoning and episodic memory retrieval. Human Brain Mapping, 37(3), 896–912. 10.1002/hbm.23074

Westphal, A. J., Chow, T. E., Ngoy, C., Zuo, X., Liao, V., Storozuk, L. A., Peters, M. A. K., Wu, A. D., & Rissman, J. (2019). Anodal Transcranial Direct Current Stimulation to the Left Rostrolateral Prefrontal Cortex Selectively Improves Source Memory Retrieval. Journal of cognitive neuroscience, 31(9), 1380–1391. 10.1162/jocn_a_01421

Yarkoni, T., Poldrack, R. A., Nichols, T. E., Van Essen, D. C., & Wager, T. D. (2011). Large-scale automated synthesis of human functional neuroimaging data. Nature methods, 8(8), 665–670. 10.1038/nmeth.1635

Zhang, Y., Brady, M., & Smith, S. (2001). Segmentation of brain MR images through a hidden Markov random field model and the expectation-maximization algorithm. IEEE Transactions on Medical Imaging, 20(1), 45–57.

