## Supplementary Materials for "What is common to memory retrieval and relational reasoning? Testing neural overlap in anterior lateral prefrontal cortex"

**DEMOGRAPHICS**

| Measure | Item | Sample | Percentage (%) |
|---|---|---|---|
| Gender | Female<br>Male<br>Non-binary | 18<br>16<br>1 | 51.4<br>45.7<br>2.9 |
| Age | 18<br>19<br>20<br>21<br>22<br>23<br>24<br>25<br>26<br>27<br>34 | 6<br>6<br>5<br>7<br>4<br>1<br>2<br>1<br>1<br>1<br>1 | 17.1<br>17.1<br>13.3<br>20<br>11.4<br>2.9<br>5.7<br>2.9<br>2.9<br>2.9<br>2.9 |
| Race | Asian<br>Asian, White<br>Black<br>Black, Pacific Islander<br>Black, White<br>White<br>White, American Indian or Alaska Native<br>Other<br>Unknown | 10<br>2<br>1<br>1<br>2<br>10<br>1<br>2<br>6 | 28.6<br>5.7<br>2.9<br>2.9<br>5.7<br>28.6<br>2.9<br>5.7<br>17.1 |
| Ethnicity | Not of Hispanic, Latino, or Spanish origin<br>Mexican, Mexican American, Chicano<br>Other Hispanic, Latino, or Spanish origin | 23<br>10<br>2 | 65.7<br>28.6<br>5.7 |
| Highest level of schooling | High school or equivalent<br>Some college, no degree<br>Associate degree<br>Bachelor’s degree<br>Master’s degree<br>Professional school degree (MD, DDS, JD) | 9<br>14<br>5<br>3<br>3<br>1 | 25.7<br>40.0<br>14.3<br>8.6<br>8.6<br>2.9 |

**Supplementary Table 1. Demographic information.**

**TASK DESIGN**

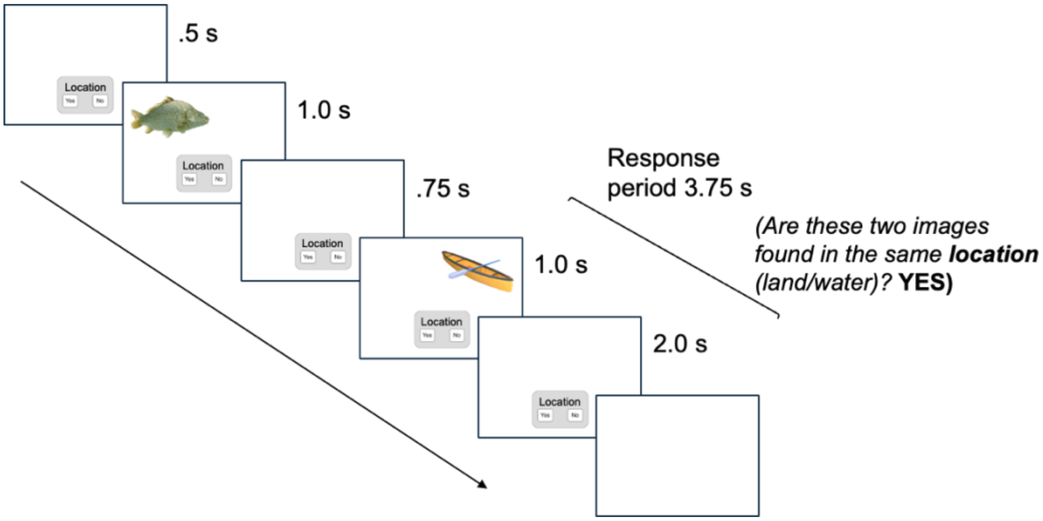


**Supplementary Figure 1. Timing of Reasoning task.** Example Vis-What 1st-order trial. A cue indicated the task-relevant dimension (in this example, Location), followed by sequential presentation of two stimuli separated by an inter-stimulus interval, and a response period. During auditory trials, a static speaker icon remained on the screen throughout stimulus presentation. Trials were separated by an intertrial interval. 1st-order trials lasted 6s. 2nd-order trials lasted 12 s and had the structure of two consecutive first-order trials with the cue "Match", and participants responded only after the second pair.

**CLUSTER PREVALENCE**

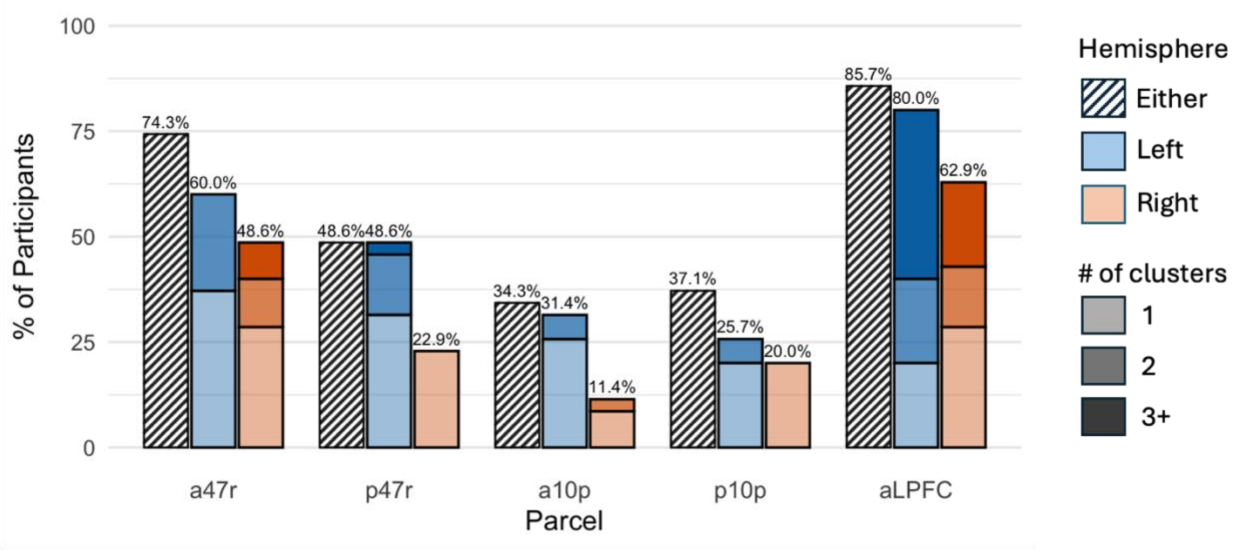


**Supplementary Figure 2. Cluster prevalence per parcel and hemisphere for the Reasoning task contrast (2nd-order > 1st-order).** Bars are stacked cumulatively, showing the % of participants with clusters at the predefined statistical threshold in the left and/or right hemispheres. Shading indicates the number of participants who had 1, 2, or 3+ clusters within a parcel.

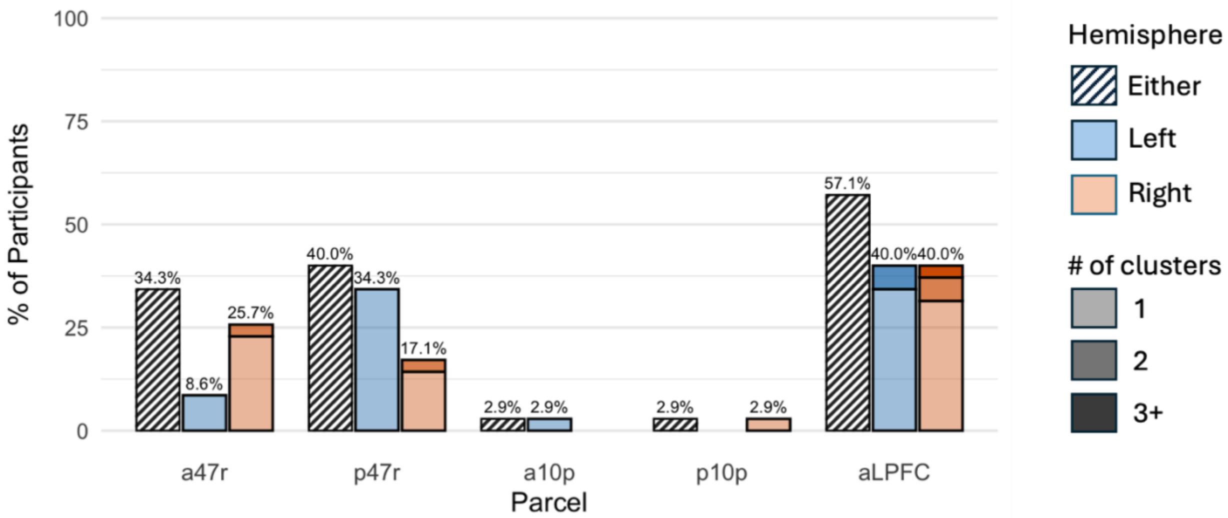


**Supplementary Figure 3. Cluster prevalence per parcel and hemisphere for the Reasoning task contrast (2nd-order > baseline).** Bars are stacked cumulatively, showing the % of participants with clusters at the predefined statistical threshold in the left and/or right hemispheres. Shading indicates the number of participants who had 1, 2, or 3+ clusters within a parcel.

**RIGHT HEMISPHERE RESULTS**

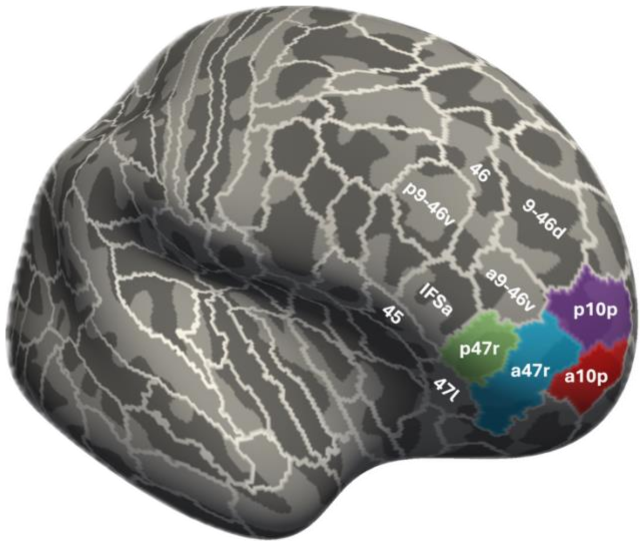


**Supplementary Figure 4. Right aLPFC parcels.** Four parcels from the Glasser multimodal cortical parcellation (Glasser et al., 2016) spanning right aLPFC. Colors indicate parcels of interest selected for analysis.

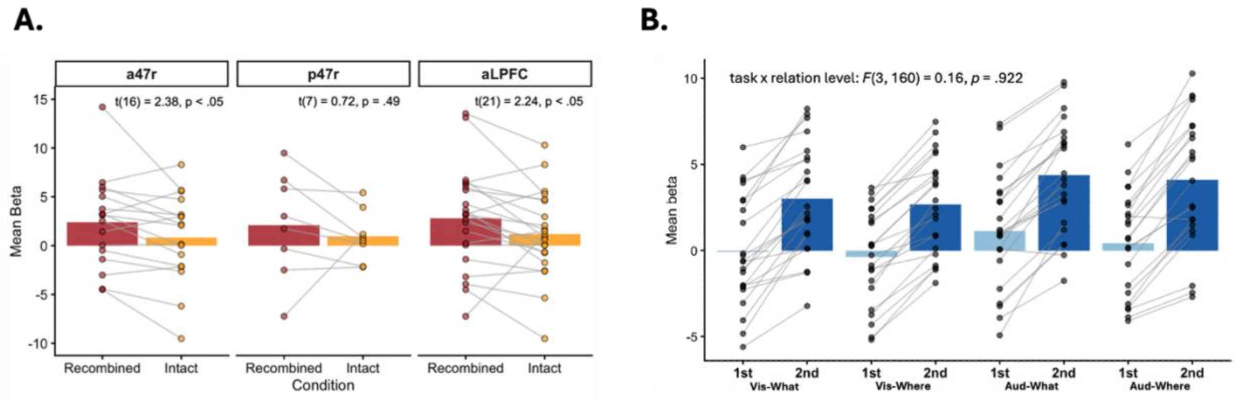


**Supplementary Figure 5. Sensitivity to memory retrieval demands and relational complexity in reasoning-related clusters in right aLPFC. (A)** Activation in reasoning-related clusters during memory retrieval. Cluster-weighted mean beta values are shown for Correct Recombined and Correct Intact trials. Points represent individual participants, and lines connect participants across conditions. Paired t-test results are shown for each parcel definition. Note that these did not survive corrections for multiple comparisons. **(B)** Sensitivity to relational complexity in reasoning-related clusters across all four Reasoning tasks. Clusters were defined using data from the entire reasoning paradigm and therefore are inherently biased; however, this verified that observed effects were not driven solely by the Vis-What task. Points represent cluster-weighted mean beta values for individual participants, and bars indicate the mean across participants.

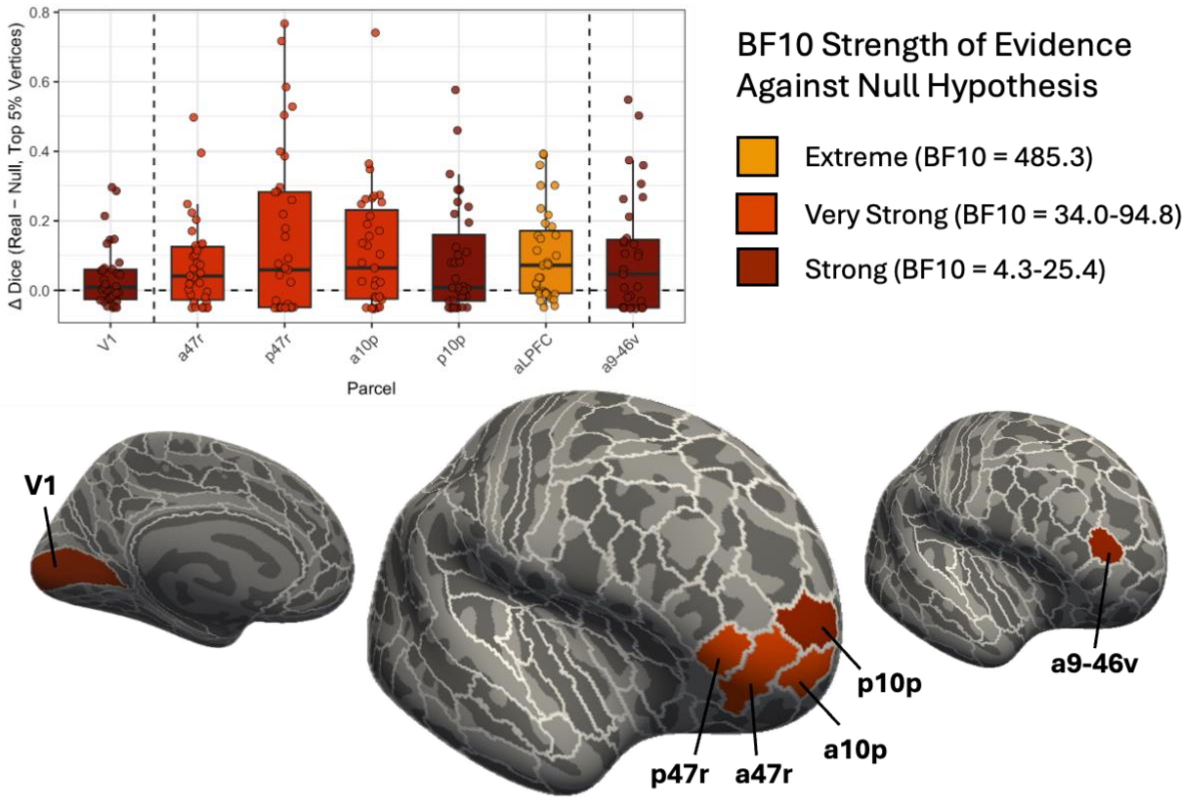


**Supplementary Figure 6. Degree of overlap between tasks varies across individuals and parcels in right aLPFC.** Colors correspond to levels of BF10 magnitude as stated in Kelter (2020). No parcels of interest showed Anecdotal or Weak levels, and thus these levels are not included here. Brains feature V1 (left), individual aLPFC parcels (middle), and a9-46v (right). The broader aLPFC parcel comprises a47r, p47r, a10p, and p10p.

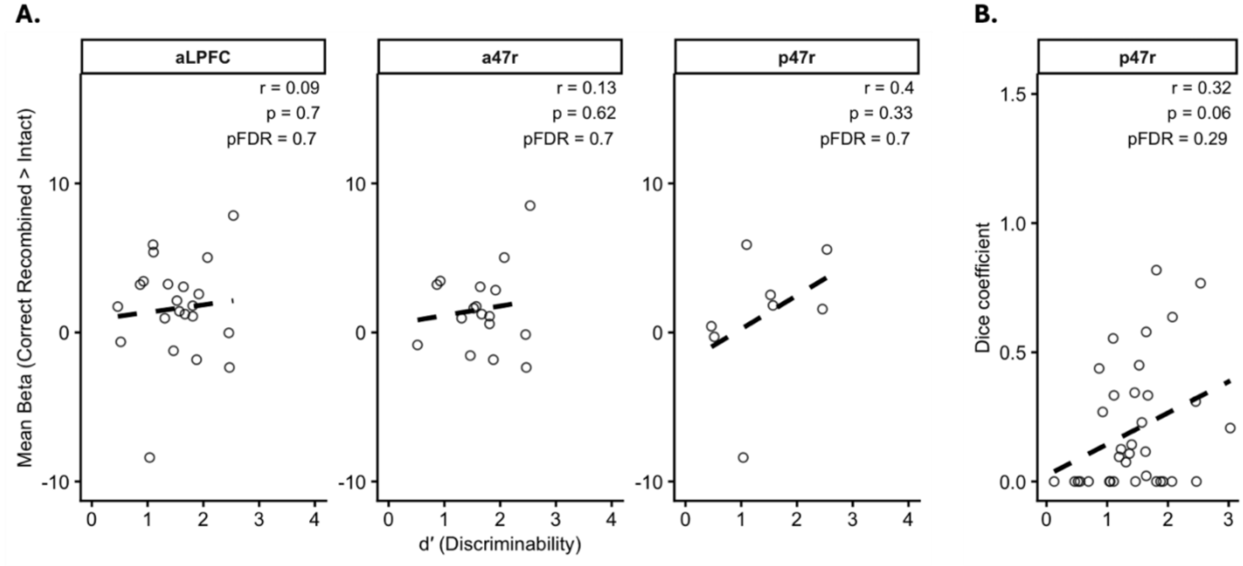


**Supplementary Figure 7. Memory performance associations with memory task activation and right aLPFC vertex-level overlap between tasks.** Each point represents one participant. Correlation coefficients (r), uncorrected p-values, and FDR-corrected p-values (pFDR) values are shown for each parcel. **(A)** Scatterplots show the relationship between recognition memory discriminability (d′) and the cluster-weighted mean beta within each parcel definition. **(B)** Scatterplot shows the relationship between recognition memory discriminability (d′) and Dice coefficients quantifying overlap between reasoning- and memory-related activation patterns (top 5% of vertices) within parcel p47r. Note that correlations between d’ and Dice coefficients were conducted for all four parcels of interest and the aLPFC parcel definition, and none were significant after FDR correction. Parcel p47r is shown here as a comparison to match the analyses in Figure 7.

**VIS-WHAT RESULTS**

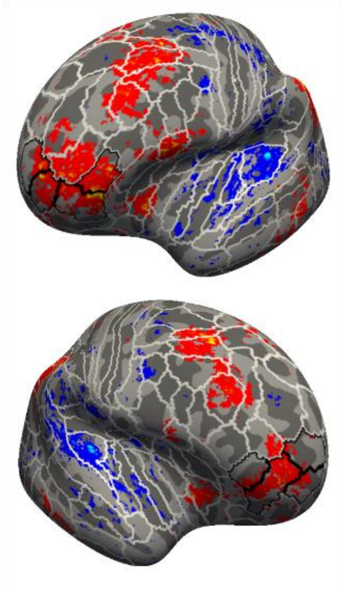

**Supplementary Figure 8. Whole-brain group-level activation contrasts during the Vis-What Reasoning task.** aLPFC parcels are outlined in black. Vis-What 2nd-order > 1st-order contrast, showing left and right hemispheres. Activation contrasts were computed with a threshold of t > 2.5.

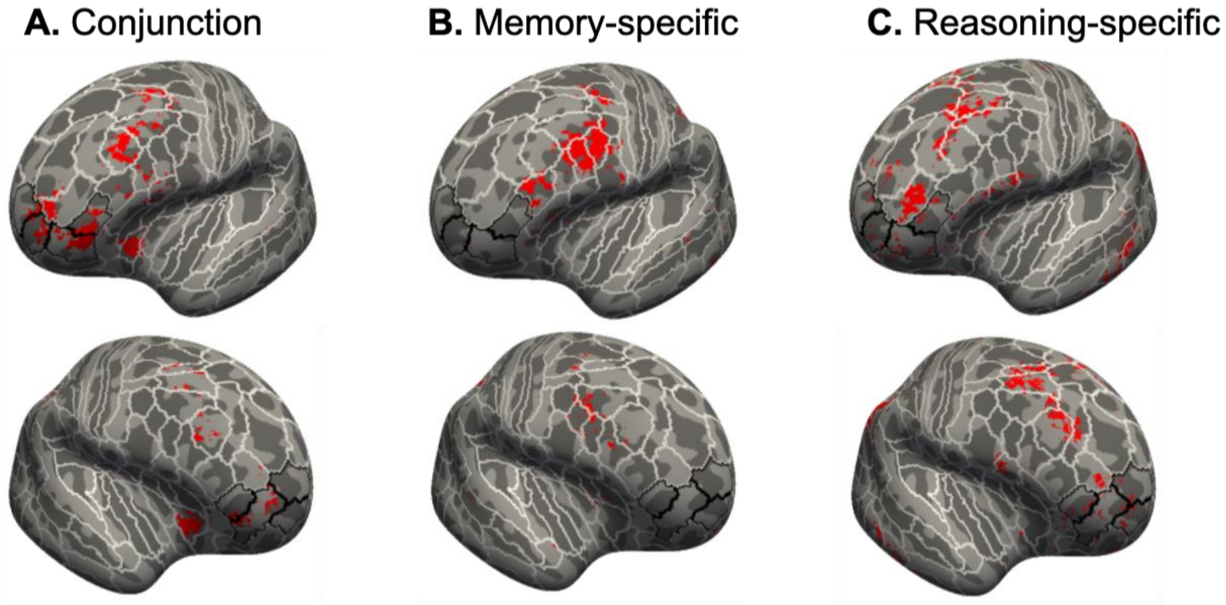


**Supplementary Figure 9. Cortical areas showing overlapping and non-overlapping activation across Memory and Vis-What reasoning tasks.** aLPFC parcels are outlined in black. **(A)** Conjunction showing overlap of activation between memory and the Vis-What Reasoning task in the left and right hemispheres, computed as the intersection of both activation contrast maps, with a threshold of $t > 2.5$. **(B)** Disjunction showing memory-specific activation. **(C)** Disjunction showing Vis-What reasoning-specific activation. Disjunction maps were computed as the intersection of task-specific activation maps, with a positive threshold of $t > 2.5$ in one task, to indicate significant activation, and a threshold of $|t| < 1$ for the other task, to indicate lack of significant activation.

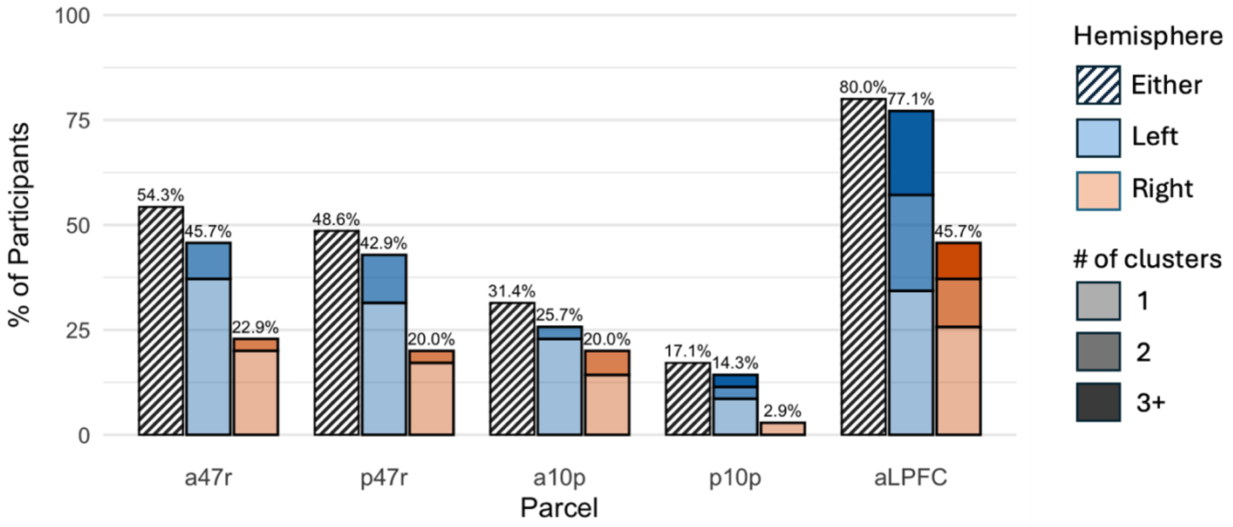


**Supplementary Figure 10. Cluster prevalence per parcel and hemisphere for the Vis-What Reasoning task contrast (2nd-order > 1st-order).** Bars are stacked cumulatively, showing the % of participants with clusters at the predefined statistical threshold in the left and/or right hemispheres. Shading indicates the number of participants who had 1, 2, or 3+ clusters within a parcel.

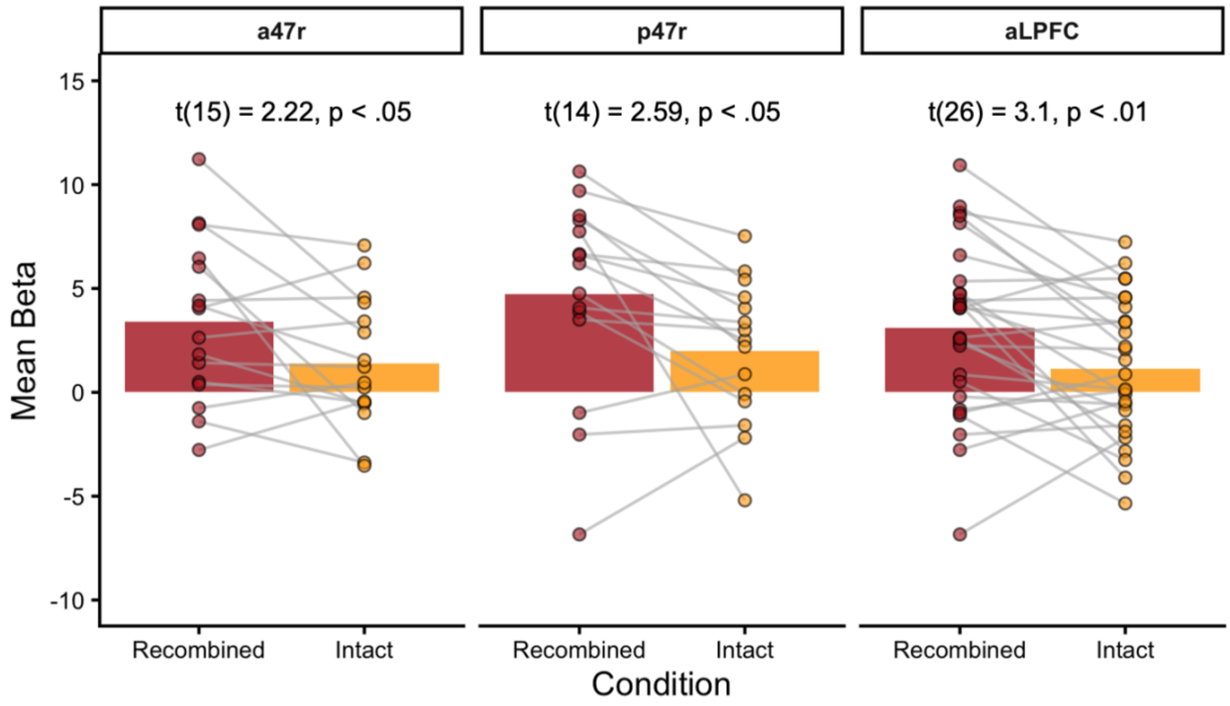


**Supplementary Figure 11. Sensitivity to memory retrieval demands in Vis-What reasoning-related clusters in left aLPFC.** Cluster-weighted mean beta values are shown for Correct Recombined and Correct Intact trials. Points represent individual participants, and lines connect participants across conditions. Paired t-test results are shown for each parcel definition.

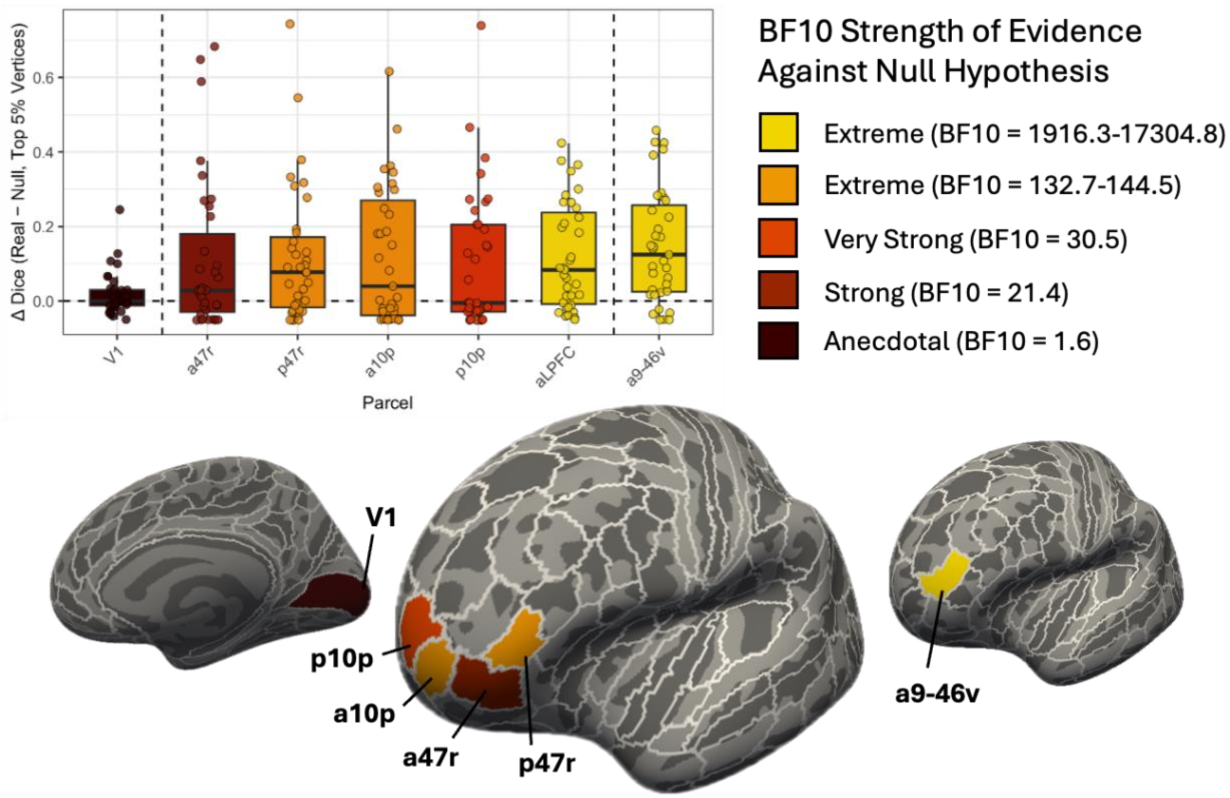


**Supplementary Figure 12. Degree of overlap between Memory and Vis-What Reasoning tasks varies across individuals and parcels in left aLPFC.** Colors correspond to levels of BF10 magnitude as stated in Kelter (2020). To emphasize differences between extremely large BF10 values, the level of "Extreme" is split into two levels. No parcels of interest showed Weak levels, and thus this level is not included here. Brains feature V1 (left), individual aLPFC parcels (middle), and a9-46v (right). The broader aLPFC parcel comprises a47r, p47r, a10p, and p10p.

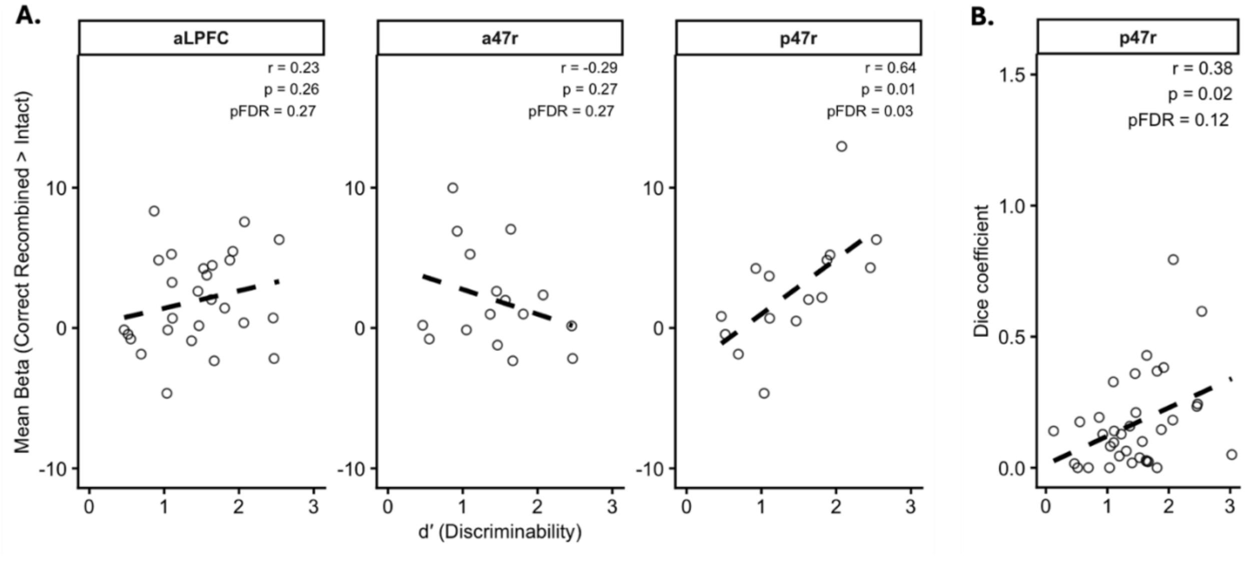


**Supplementary Figure 13. Memory performance associations with memory task activation and vertex-level overlap between Memory task and Vis-What Reasoning task.** Each point represents one participant. Correlation coefficients (r), uncorrected p-values, and FDR-corrected p-values (pFDR) values are shown for each parcel. **(A)** Scatterplots show the relationship between recognition memory discriminability (d′) and the Vis-What reasoning cluster-weighted mean beta within each parcel definition. **(B)** Scatterplot shows the relationship between recognition memory discriminability (d′) and Dice coefficients quantifying overlap between Vis-What reasoning- and memory-related activation patterns (top 5% of vertices) within parcel p47r. Note that correlations between d’ and Dice coefficients were conducted for all four parcels of interest and the aLPFC parcel definition, and none were significant after FDR correction. Parcel p47r, shown here, was the only significant parcel before correction.
